# Beyond organ age gaps: comparing plasma proteomic metrics of organ biology in relation to brain structure and neurodegenerative disease

**DOI:** 10.64898/2026.11.09.765339

**Authors:** Romina Zendehel, Lijun An, Léa Chauveau, Hamilton Se-Hwee Oh, Gabriele Vilkaite, Ines Hristovska, Varun Warrier, Jakob Seidlitz, Richard A.I. Bethlehem, Alexa Pichet-Binette, Jacob W. Vogel

## Abstract

Biological aging is heterogeneous across organ systems, and brain health may be influenced by biological processes extending beyond the central nervous system (CNS). This can be investigated using biological age measures that are increasingly used to estimate how biological systems deviate from chronological aging. However, organ level plasma proteomic metrics may capture different forms of biological variation depending on how they are defined. Here, we compared three complementary organ level proteomic metrics across 15 organ systems (and two global systems) in relation to age, organ-relevant physiological phenotypes, brain morphometry, and incident neurodegenerative disease, using plasma proteomics from 52,667 UK Biobank participants and brain MRI data from 5,078 of those individuals. We derived i) supervised age gap metrics using XGBoost models trained to predict chronological age, ii) unsupervised PC1 metrics capturing dominant organ-associated proteomic variation, and iii) phenotype guided PLS metrics linking plasma proteins to organ-relevant physiological traits. The three metric classes were related but not interchangeable. Global all-protein and all-organ-protein models showed strong age-related proteomic structure. However, organ-specific PC analyses showed that age was often but not always the dominant source of variation in protein signal across organs. Age gap metrics showed the most consistent associations with subsequent ICD-10based neurodegenerative and vascular diagnoses, whereas all three metric classes showed distinct associations with brain structure and organ-relevant physiological phenotypes. Brain structure analyses revealed heterogeneous organ–brain association patterns across organs, suggesting that proteomic variation relates to brain structure in both a systemic and an organ-specific manner. Several systemic and organ-specific signals partially mediated the associations between disease specific genetic risk and incident diagnoses of Alzheimer’s disease and stroke, albeit with small effect sizes (1-8%). Similarly, organ metrics provided modest incremental predictive improvement over baseline demographic models, with the largest increase in ROC-AUC observed for Alzheimer’s disease when all organ age gap metrics were added jointly (ΔROC-AUC = 0.037). These results link brain related outcomes to proteomic variation across both CNS and non-CNS systems. Our work suggests that proteomic organ metrics capture complementary dimensions of biological variation rather than a single construct of organ aging. Although these measures may provide informative biological and potentially predictive signals, their incremental predictive value in the present analyses was modest.

## Introduction

Biological aging is a systemic and heterogeneous process involving multiple organ systems that may age at different rates within the same individual^1–3^. Age related molecular and functional changes do not progress uniformly; instead, they accumulate at various rates across different tissues. This heterogeneity holds biological significance, as the accelerated aging of specific organs correlates with a heightened risk of organ-specific diseases and mortality^4^. At the same time, organ aging is not an isolated process; evidence suggests that dysfunction in peripheral systems, especially vascular, immune, and metabolic systems, plays a significant role in the health or dysfunction of the brain^4^.

In this context, neurodegeneration may partly reflect the cumulative burden of age-related dysfunction across organ systems, potentially increasing the vulnerability of the aging brain^5,6^. While the brain exhibits unique hallmarks of decline, such as bioenergetic failure, structural atrophy, and the aberrant cell cycle re-entry of post mitotic neurons^7–9^, these processes do not occur in isolation. Neuroimaging-derived “brain age” has emerged as a marker of brain structural aging^10^. In addition, growing evidence suggests that biological variation outside the CNS is associated with brain health^10^. Peripheral organ dysfunction may influence brain health through pathways involving chronic inflammation, blood-brain barrier damage, and impeded peripheral clearance of neurotoxic proteins, which have been implicated in Alzheimer’s and Parkinson’s diseases^6,11^. Age related vascular dysfunction has been associated with cognitive decline and neurodegenerative processes, potentially through impaired cerebral perfusion and neurovascular integrity, emphasizing the role of extra neural processes in shaping brain aging trajectories^12–14^.

One approach to fully capture the crosstalk between the brain and the body is to study molecular dynamics occurring within the circulatory system. Circulating plasma proteins offer an integrative perspective on organism wide physiology by reflecting both local tissue conditions and systemic biological regulation^3,15^. Large-scale plasma proteomic profiling combined with machine learning approaches has enabled the development of organ-specific aging models based on tissue associated circulating proteins, offering a minimally invasive approach to estimate biological age across organ systems^1,16,17^. The organ age gap, characterized as the difference between predicted and chronological age, is commonly interpreted as a measure of accelerated or decelerated organ aging^1,2^. In large population studies, organ-specific age gaps show low to moderate correlations across organ systems, consistent with distinct patterns of biological aging with individuals^1,18–20^. Higher organ age gaps have also been associated with organ-specific disease, mortality, and other health outcomes^1,2,16,21^. Age gap-based organ metrics are optimized to capture proteomic variation associated with chronological age. Yet proteomic signatures of organ integrity may also reflect functional decline or dysregulation that is partially or fully independent of age. It therefore remains unclear how this age anchored dimension relates to other biologically relevant dimensions of organ level proteomic variation.

This study systematically compares several complementary organ level proteomics metrics across 15 organ systems, assessing their relationships with organ-specific physiological phenotypes, and investigating how they relate to brain structure and neurological disease. Specifically, in addition to organ age gaps, we investigate proteomic based organ function metrics that are anchored to organ-specific phenotypes, or to dominant organ-specific variance components. By comparing these complementary representations, we aimed to understand the influence of peripheral organs on brain health, and to do so while disentangling age-related from non-related aspects of organ function.

## Methods

### Study Overview and Objectives

This cross-sectional observational study analyzed organ-specific biological aging and neurodegenerative features using high dimensional plasma proteomics, clinical biomarkers, and neuroimaging data. The main objective was to characterize organ-specific functional and physiological states using circulating protein signatures. These metrics were compared with brain structure, future clinical diagnoses, and genetic risk for neurodegenerative disease to evaluate how systemic biological variation relates to brain health and disease vulnerability (Figure 1).

**Figure 1.**
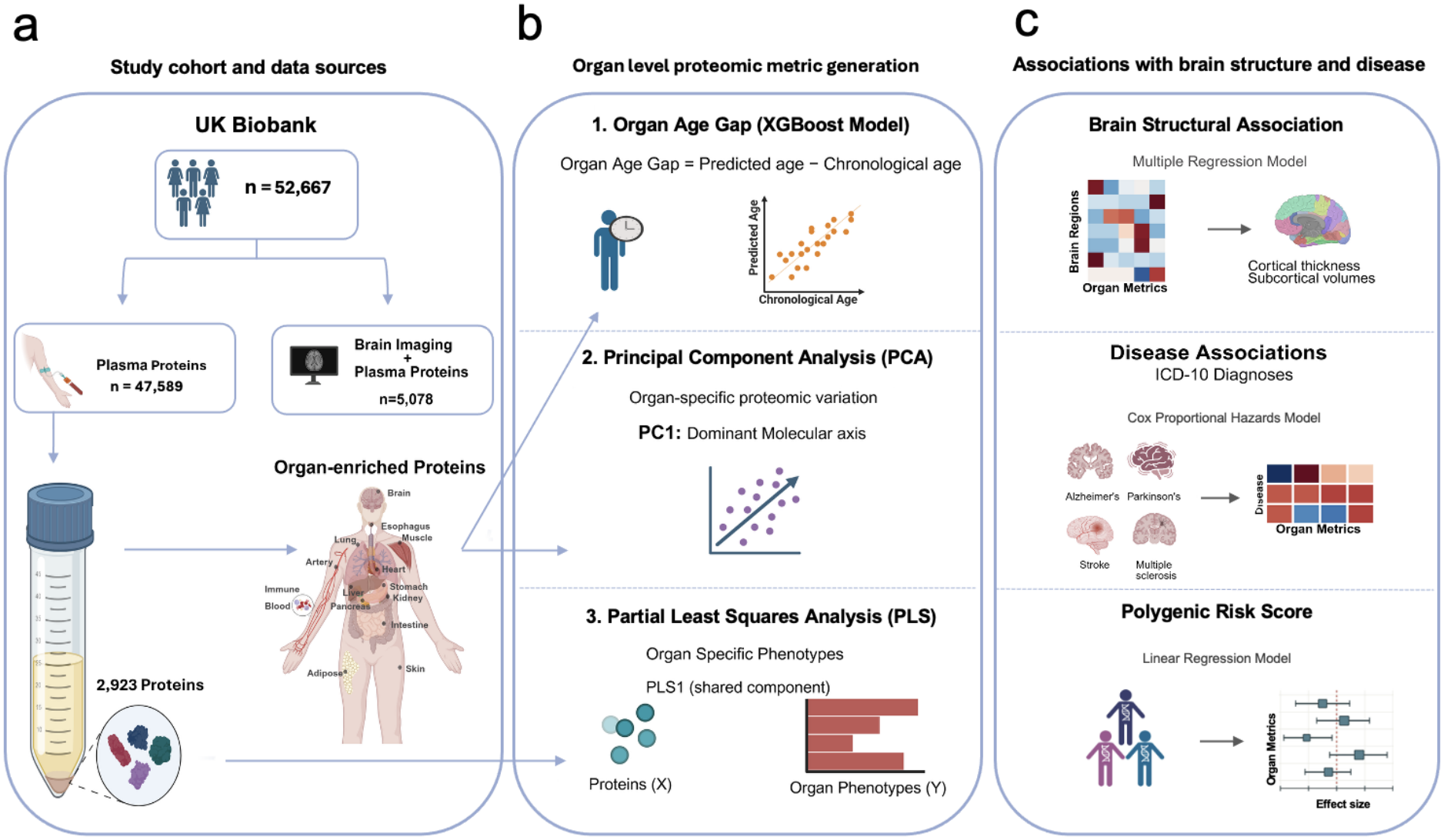
Schematic overview of the study design and analytical framework. **a)** Study cohort and data sources. Plasma proteomics (n= 52,667) and neuroimaging data (n= 5,078) from the UK Biobank were analyzed to create organ-specific protein panels for 15 organ systems. **b)** Organ level proteomic metric generation. Three complementary proteomic health metrics were produced: (1) the organ age gap estimated using extreme gradient boosting (XGBoost), (2) the dominant axes of proteomic variation identified through principal component analysis (PCA; PC1), and (3) phenotype informed components obtained from partial least squares regression (PLS; PLS1). **c)** Association with brain structure and disease. Organ level measures were examined for associations with brain structure (cortical thickness and subcortical volumes) using multiple regression models, incident ICD-10, defined clinical diagnoses by Cox proportional hazards models, and polygenic risk scores for neurodegenerative using linear regression. This figure was created with BioRender.com.

### Study Cohort

This study used data from the UK Biobank, a population-based cohort of more than 500,000 participants aged 40 - 69 years at recruitment between 2006 and 2010. At baseline, participants underwent physical assessments, provided biological samples, and completed questionnaires on health and lifestyle. UK Biobank obtained ethical approval from the Northwest Multi-Centre Research Ethics Committee (REC reference: 16/NW/0274).

The main analytical cohort included 52,667 participants with plasma proteomic data (Table 1). Baseline was defined as UK Biobank recruitment. For supervised organ age prediction, two analytical cohorts were established from this subset: (i) a model development cohort comprising 47,589 participants with proteomics data but without neuroimaging, used for training and validation of supervised biological age models; and (ii) an independent imaging test cohort comprising 5,078 participants with both proteomics and brain MRI data, used for independent model evaluation and organ–brain association analyses. Plasma samples were collected at baseline recruitment, while brain MRI were performed at later UKB imaging visits (instances 2 and 3), therefore the two measures were not taken at the same time. After model development, predicted biological ages and values were generated for the broader proteomics cohort and used in downstream analyses.

**Table 1.** Cohort Structure, Dataset Allocation, and Baseline Characteristics.

| **Cohort Structure and Dataset Allocation** |  |  |  |  |  |
|  | **Data subset** | **N** | **Age, years (mean ± SD)** | **Female sex, n (%)** |  |
|---|---|---|---|---|---|
| **Full proteomics cohort** | Plasma proteomics | 52,667 | 56.8 ± 8.2 | 28,395 (53.9) |  |
| **Model development set** | Proteomics only | 47,589 | 57.1 ± 8.2 | 25,748 (54.1) |  |
| Internal training subset | Proteomics only | 38,071 | 57.1 ± 8.2 | 20,608 (54.1) |  |
| Internal validation subset | Proteomics only | 9,518 | 57.1 ± 8.2 | 5,140(54.0) |  |
| **Independent imaging test set** | Proteomics +<br>brain MRI | 5,078 | 54.4 ± 7.8 | 2,647 (52.1) |  |
| **Incident Neurological Diagnoses: Baseline Characteristics** |  |  |  |  |  |
|  | **Alzheimer’s<br>(n = 699)** | **Parkinson’s<br>(n = 884)** | **MS<br>(n = 234)** | **Stroke<br>(n = 454)** | **ALS<br>(n = 286)** |
| **Age, years (mean ± SD)** | 64.6 ± 5.3 | 63.2 ± 5.1 | 54.6 ± 7.8 | 62.2 ± 6.1 | 60.3 ± 6.8 |
| **Female sex (%)** | 45.9 | 62.6 | 30.3 | 55.9 | 54.9 |

Different downstream analyses used the maximum available overlap between proteomic metrics and the relevant phenotype, imaging, diagnosis, or genetic data. PCA - derived metrics were calculated in the full proteomics cohort. PLS-derived metrics were trained in organ-specific phenotype complete subsets. The fitted PLS protein weights were then applied to all participants with available preprocessed plasma proteomics data to generate participant level PLS1 scores. Brain structure analyses were performed in the imaging subset. Clinical diagnosis analyses were performed in participants with both available organ level proteomic metrics and ICD-10 linked follow-up data. PRS analyses were performed in the overlap between available polygenic risk scores and proteomic metrics.

Plasma proteomics data were obtained from the UK Biobank Pharma Proteomics Project (UKB-PPP). Protein levels were quantified using the antibody based Olink Explore 3072 proximity extension assay platform, which measured 2,923 analytes in 54,219 participants before downstream filtering^22^. Quality control procedures were implemented centrally by UKB-PPP prior to data release. These included removal of Olink control samples, exclusion of withdrawn or unprocessed participants, outlier sample detection using principal component analysis and IQR-median approaches, and filtering of measurements flagged with quality control or assay warnings. The released dataset consisted of quality controlled normalized protein expression (NPX) values.

For the present analyses, the final proteomics dataset included 2,923 proteins in 52,667 participants. Additional preprocessing was applied before organ level modeling. Protein level outliers were identified using a z-score threshold of |z| > 5 and masked as missing values. Remaining missing values were imputed using feature wise mean imputation. Following imputation, protein variables were standardized to zero mean and unit variance before downstream analyses. Global proteomic PC1 residualization was applied, where the first principal component was extracted from the standardized protein matrix and regressed out to reduce dominant shared variation across proteins^23^.

#### Organ-specific protein mapping

For each of the 15 organs, curated lists of tissue enriched proteins^1,24^ were used to construct organ-specific input matrices. These organ enriched protein sets were originally defined using GTEx and Human Protein Atlas-derived transcriptomic criteria, in which genes expressed at least four-fold higher in one organ compared with all other organs were classified as organ enriched. Gene level organ annotations from Oh et al^1^. were mapped to the measured Olink plasma proteome to generate organ-specific protein panels for adipose, artery, brain, esophagus, heart, immune system, intestine, kidney, liver, lung, muscle, pancreas, skin, stomach, and whole blood. The full organ protein mapping, including the number of mapped proteins per organ, is provided in (Supplementary Table 1). Two global protein sets were also generated. The organ-specific proteins model included the union of all proteins present in both the proteomics matrix and the organ protein mapping. The all-proteins model included all available plasma proteins in the final proteomics matrix.

#### Phenotypic, clinical, and neuroimaging data

Quantitative phenotypes and clinical diagnosis data were obtained from UK Biobank field data, imaging-derived phenotypes, and linked hospital inpatient records^25,26^. organ-relevant phenotypes were selected based on biological relevance, quantitative availability, and their relationship to organ structure or function. These phenotypes were grouped by organ system and are listed in (Supplementary Table 2).

Clinical disease outcomes were defined using ICD-10 codes from linked health records^26^. Neurodegenerative and related diagnoses were encoded as binary outcomes, where participants were classified as cases if they had at least one qualifying ICD-10 code for the disease of interest during follow-up. Diagnoses occurring before baseline were excluded from incident disease analyses. Follow-up time was calculated as the time from baseline to the first qualifying diagnosis or censoring. The exact ICD-10 codes used for each disease category are provided in (Supplementary Table 3).

Structural MRI data were obtained from the UK Biobank imaging extension^27^. Imaging analyses included 5,078 participants with both plasma proteomics and brain MRI data. Neuroimaging features included regional cortical thickness and subcortical volumes generated by the UK Biobank imaging pipeline (image-derived phenotypes; IDPs)^28,29^. Cortical thickness measures were derived using FreeSurfer-based surface reconstruction and parcellated according to the Desikan–Killiany atlas^29,30^. Thirty-one bilaterally averaged regional cortical thickness measures were included in organ cortical thickness association analyses. Subcortical structures were segmented using FMRIB’s Integrated Registration and Segmentation Tool, and 12 subcortical volume measures were included^29,31^.

For PLS modeling (see below), cortical thickness regions were excluded from the brain phenotype set to avoid circularity, as cortical thickness was later used as an outcome in organ– brain association analyses. Therefore, the neuroimaging feature set used for brain PLS models consisted of other brain morphometrics (Supplementary Table 2). Imaging features were preprocessed using the same general framework as other quantitative variables, including outlier masking, mean imputation, and z-score normalization. For subcortical volume association analyses, total intracranial volume (TIV) was included as a covariate in downstream statistical models.

#### Derivation of organ level proteomic metrics

To characterize complementary dimensions of organ related proteomic variation, three organ level proteomic metrics were derived: age gap, PC1, and PLS1. Age gap was defined as an age anchored supervised metric, PC1 as an unsupervised metric capturing the dominant axis of organ-associated proteomic variation, and PLS1 as a phenotype guided metric capturing proteomic variation related to organ-relevant phenotypes (Figure 1).

### Supervised organ age prediction and age gap calculation

Supervised organ age models were trained to estimate chronological age from plasma proteomic profiles, following the general framework of proteomic organ age modelling and age gap estimation^1,24^. Separate models were trained for each of the 15 organ-specific protein panels defined by the protein to organ mapping: adipose, artery, brain, esophagus, heart, immune system, intestine, kidney, liver, lung, muscle, pancreas, skin, stomach, and whole blood. In addition to these organ-specific models, two global age prediction models were trained to capture broader proteomics signals; an organ-specific proteins model using the union of all organ enriched proteins, and all proteins model using the full available plasma proteomic feature set. For each organ-specific model, input features included the corresponding organ-specific plasma proteins and sex. No additional covariates were included in the supervised age prediction models.

Chronological age at recruitment, in years, was used as the prediction target. Models therefore predicted raw chronological age rather than normalized, scaled, residualized, or transformed age. Extreme gradient boosting regression was implemented using XGBoost with squared error loss^32^. Participants were split according to imaging availability: participants without imaging data were used for model development, and participants with imaging data were held out as an independent test set. The model development set included 47,589 participants and was further divided into internal training and validation subsets using an 80/20 split (Table 1). The independent imaging test set included 5,078 participants.

Hyperparameters were optimized using GridSearchCV with five-fold cross-validation and R² as the scoring metric. The hyperparameter grid included maximum tree depth values of 3, 5, and 7; numbers of estimators of 100, 200, 300, and 500; and learning rates of 0.01, 0.05, and 0.1. After hyperparameter selection, the best model was used to generate age predictions in the training, validation, and independent test sets. Model performance was evaluated using R², Pearson correlation, mean absolute error, and root mean squared error, with the independent test set evaluated once after model development (Supplementary Table 4).

For each participant, raw age gap was calculated as predicted biological age minus chronological age:

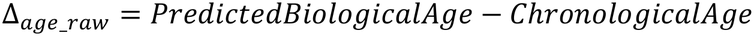

Positive raw age gap values indicated older predicted biological age relative to chronological age, whereas negative values indicated younger predicted biological age. To reduce dependence of age gap on chronological age, raw age gap was residualized against chronological age and an orthogonalized quadratic age term. Specifically, age² was first orthogonalized with respect to age, and raw age gap was then regressed on age and the orthogonalized age² term. The residual from this model was retained as the corrected age gap for downstream analyses^14^. Positive corrected age gap values indicate relatively older organ biological age after accounting for chronological age, while negative values indicate relatively younger organ biological age. Model calibration was visualized by plotting predicted versus chronological age in the independent test set (Supplementary Figure 1). We acknowledge that regressing age and age derivatives from predicted age gap muddles interpretability of predicted age gap. However, without this step, all age gaps are nearly collinear. We view this issue as a built-in limitation of the age gap approach for organ aging.

### Principal component analysis

Principal component analysis was used to derive an unsupervised organ level proteomic metric^33^. Organ age gap approaches estimated age-related deviation by training models to predict chronological age^1^, but this assumes that the dominant biologically relevant signal in an organ-specific proteomic panel is age-related. To test this assumption, we used PCA to summarize the dominant axis of coordinated protein variation within each organ-specific protein set without using chronological age as model target. This allowed us to evaluate whether the major source of organ-associated proteomic variation was aligned with aging or instead reflected other biological variation. PCA was performed separately for each organ-specific protein set using proteins assigned to that organ by the protein to organ mapping and present in the proteomics dataset (Supplemental Table 1). PCA was performed in the full proteomics cohort, and the transforms applied directly to the independent imaging test set.

PCA was applied to the preprocessed organ-specific protein matrices. For each organ, the first three principal components were extracted. PC1 scores were used as the main unsupervised organ level proteomic metric in downstream analyses, representing the dominant axis of organ-associated proteomic variation. PC2 and PC3 were retained for supplementary analyses examining whether age-related protein variation aligned with higher order axes of proteomic variation (see below).

PC loadings from PC1, PC2, and PC3 together with explained variance and protein contribution values, were saved and used to evaluate the relationship between PCA-derived components and protein–age associations (Supplementary Table 5). To provide a consistent direction across organs, PC1 scores and loadings were multiplied by −1 when the PC1 score was negatively correlated with chronological age. Thus, higher PC1 values were oriented to reflect an older like proteomic profile, and positive PC1 loadings indicated proteins contributing to this older like direction. PC1 was not assumed to represent organ dysfunction, but rather population variation in organ-specific proteins. PCA was implemented using sklearn. decomposition PCA in scikit-learn34.

### Partial least squares regression

Partial least squares regression was used to derive phenotype guided organ level proteomic metrics^35^. PLS models were implemented using PLSRegression in scikit-learn with one latent component retained for downstream analyses^34^.

A separate PLS model was fitted for each organ with available organ-relevant phenotype data. In contrast to the age gap and PCA approaches, which summarized variation within organ-specific protein panels, PLS was used to identify proteomics patterns that covaried with quantitative indices of organ structure or function. Because these phenotype linked signals may involve systemic proteins not classified as organ enriched, PLS was used on all available plasma proteins as predictor variables rather than only the organ enriched protein panel. This provided a complementary phenotype guided metric for downstream comparisons with the other organ level proteomic metrics. Specifically, the full plasma proteomic feature matrix was used as X, while the Y matrix consisted of manually selected organ-relevant quantitative UK Biobank phenotypes. Phenotypes were selected based on organ relevance, quantitative availability, sufficient non-missing data, and interpretable links to organ structure or function. The selected phenotypes and their UK Biobank field IDs are listed in Supplementary Table 2.

Before PLS fitting, both the protein matrix and phenotype matrix were residualized for sex and height. These covariates were applied consistently across organ models to reduce variation attributable to sex and body size, which can strongly influence several physiological phenotypes, particularly musculoskeletal and body composition related measures. Organ phenotypes were not imputed participants were included only if they had complete organ phenotype and covariate data. Protein values were taken from the preprocessed proteomics matrix described above. The PLS implementation in scikit-learn was then used with its default internal scaling of X and Y. Age was not included as a covariate in the PLS models.

Participant level PLS1 scores were extracted from the X side of the model using the residualized protein matrix. Protein weights and phenotype weights were retained for downstream interpretation of the phenotype guided components. PLS scores were oriented using predefined phenotype direction anchors so that higher PLS1 values corresponded to the less favorable direction of the phenotype guided profile. Because the sign of a PLS component is arbitrary, this orientation was used only to facilitate biological interpretation. For whole blood, the initial literature informed orientation was inconsistent with the dominant phenotype weight pattern and with the broader direction of the PLS signals across analyses. The whole blood PLS score was therefore reversed before downstream analysis PLS1 protein loadings were inspected to describe molecular contributors to phenotype guided organ axes (Supplementary Figure 3). Models were defined for organs with at least three organ-relevant phenotype data, including brain, lung, pancreas, heart, artery, kidney, liver, muscle, adipose, immune system, whole blood, and skin (Supplementary Table 6).

Because PLS1 was derived using organ-relevant phenotypes, subsequent associations between PLS1 and those phenotypes were interpreted as phenotype guided specificity rather than as independent validation of metric performance.

### Statistical analyses

#### Correlations among organ level proteomic metrics

Pairwise Pearson correlations among organ level proteomic metrics were calculated to characterize the correlation structure across age gap, PC1, and PLS1. For age gap and PC1, analyses additionally included the all organ-specific protein model and all-protein model. The final correlation matrix was calculated across with complete data. No p-values were calculated for this analysis, which was interpreted descriptively to visualize the correlation structure among organ level proteomic metrics.

#### Alignment between PCA loadings and protein–age associations

To assess whether PCA-derived components reflected age-related protein variation, protein–age associations were first calculated for each protein as Pearson correlations between protein abundance and chronological age. These protein age correlations were calculated without covariate adjustment. For each organ-specific protein set, PC1, PC2, and PC3 loadings were extracted from the PCA models. Spearman correlations were then calculated between protein loadings and the corresponding protein age Pearson correlations within each organ and principal component.

This analysis included 15 organ-specific protein groups and two global protein groups: the organismal/all organ-specific protein model and the all-proteins model. In total, 51 group by component tests were performed. Benjamini–Hochberg false discovery rate (FDR) correction was applied across all 51 tests. Adjusted p-values are reported in Supplementary Table 7.

#### Associations with physiological phenotypes

Associations between organ level proteomic metrics and physiological phenotypes were assessed using pairwise Pearson correlations. Phenotypes were grouped by organ assignment using the organ labels in the phenotype metric association table (Supplementary Table 2).

To assess phenotype specificity, phenotypes assigned to the same organ as the proteomic metric were treated as organ matched phenotypes, while phenotypes assigned to other organs were treated as non-matched phenotypes. Phenotype specificity was summarized as the ratio of the mean absolute Pearson correlation with organ matched phenotypes to the mean absolute Pearson correlation with non-matched phenotypes. A specificity ratio greater than one indicated stronger associations with phenotypes from the corresponding organ than with phenotypes from other organ systems.

No covariate adjustment or FDR correction was applied in this exploratory phenotype association analysis. Because PLS1 metrics were derived using organ-relevant phenotype matrices, associations between PLS1 and those phenotypes were interpreted as phenotype guided specificity rather than independent validation of metric performance.

#### Associations with brain structure

Brain structure associations were tested in the imaging subset of participants with both plasma proteomics and brain MRI data (n = 5,078). Proteomics sampling and MRI acquisition occurred at different UKB assessment visits and were therefore non-concurrent. Outcomes included 31 regional cortical thickness measures derived from the Desikan–Killiany atlas and 12 subcortical volume measures derived from aseg-style segmentations. For both cortical thickness and subcortical volumes, corresponding left and right hemisphere measures were averaged to obtain bilateral values. Age gap and PC1 analyses included both cortical thickness and subcortical volume outcomes, whereas PLS1 analyses included cortical thickness outcomes only; subcortical PLS1 associations were not modeled because subcortical measures were included among the phenotypes used to derive the brain PLS score and were therefore not evaluated again as downstream outcomes to avoid circularity.

For each brain structural outcome, ordinary least squares regression models were fitted with organ level proteomic metrics as candidate predictors. Models were run separately for each metric family: age gap, PC1, and PLS1. Within each family, all retained organ level proteomic metrics were included together in the same model to estimate organ-specific associations while accounting for shared variance among correlated organ metrics. Prior to this step, multicollinearity among candidate organ level proteomic predictors was assessed using variance inflation factors (VIFs), as overlapping or related proteomic data can show redundancy, leading to unstable coefficient estimates when included jointly. A VIF threshold of 5 was used to identify problematic multicollinearity^36^. For age gap models, no organ level predictors exceeded this threshold, and all organ-specific age gap metrics were retained as candidates. For PC1 models, the all-proteins PC1 metric showed high collinearity with other PC1-derived metrics and was excluded from the candidate set, while all the remaining PC1 metrics were retained. For PLS1 models, stronger collinearity was observed across organ level PLS1 scores, consistent with the use of all available plasma proteins as predictors in the PLS models. Based on VIFs only brain, immune, and skin PLS1 scores were retained. Exclusion from multivariable models reflected statistical redundancy among predictors and did not imply that excluded organ metrics were biologically irrelevant. Age and sex were included as covariates in all models. For subcortical volume models, eTIV was additionally included as a covariate. Imaging assessment center was additionally included as a categorical covariate in all brain structure models to account for site related variation in MRI measurements. Participants were scanned at Cheadle, Newcastle, reading, or Bristol. Because only four participants in the analytic sample were scanned at Bristol, these participants were excluded, leaving 5,074 participants across the three remaining imaging centers. Imaging center indicators were retained as prespecified covariates in all models.

Candidate organ level proteomic metrics were selected using AIC-based bidirectional stepwise selection. Models started with prespecified covariates, and candidate organ metrics were iteratively added if they reduced AIC. Candidate organ predictors were then removed if their removal further reduced AIC. This procedure was repeated until no additional AIC improvement was observed. Prespecified covariates were retained in all models. Therefore, organ metrics were selected based on AIC improvement rather than p-value entry or removal thresholds. For each retained organ metric, model coefficients, t-values, and p-values were extracted from the final OLS model. To complement these multivariable models, single metric analyses were performed, in which each organ level proteomic metric was tested separately for its association with each brain structural outcome, using covariate adjusted OLS models. These models were adjusted for age and sex, imaging center, with eTIV additionally included for subcortical volume outcomes.

Benjamini–Hochberg FDR correction was applied separately within each metric family result table, using the non-null p-values from retained associations. For the main bidirectional models, FDR correction was therefore applied separately for age gap, PC1, and PLS1 results, rather than pooled across all metric families or all possible unselected organ region combinations.

#### Associations with incident clinical diagnoses

Associations between baseline organ level proteomic metrics and incident diagnoses of Alzheimer’s disease, Parkinson’s disease, amyotrophic lateral sclerosis, multiple sclerosis, and stroke were tested using ICD-10 based disease definitions. Cause specific Cox proportional hazards models were fitted separately for each diagnosis and each proteomic metric family; age gap, PC1 and PLS.

Baseline was defined as the initial UK Biobank assessment visit. For each disease, participants with a qualifying diagnosis occurring before or at baseline were excluded. Follow-up time was calculated from baseline to the earliest of incident diagnosis, death, administrative end and follow-up, and/or earlier documented end of follow-up, where applicable. Participants without an incident diagnosis during follow-up were censored at their corresponding end of follow-up date. Participants with missing baseline or censoring information, unknown diagnosis timing, or invalid follow-up intervals were excluded.

Cox models were adjusted for age and sex, with follow-up time incorporated as the survival time rather than as a model covariate. Single metric models were fitted separately for each organ level proteomic metric. For multivariable analyses, candidate proteomic metrics were entered using the same VIF-based candidate selection procedure described above, followed by bidirectional AIC-based stepwise selection. Age and sex were retained as prespecified covariates throughout model selection.

For each retained term, hazard ratios (HRs), 95% confidence intervals, Wald statistics, and p-values were extracted from the final Cox model. Benjamini–Hochberg FDR correction was applied separately within each metric family result table. The proportional hazards assumption was assessed using Schoenfeld residual based tests, and model diagnostics were reviewed for violations of this assumption.

#### Polygenic risk score analyses

Polygenic risk score (PRS) analyses were performed to assess whether organ level proteomic metrics were associated with genetic liability for neurodegenerative and vascular disease. PRS for Alzheimer’s disease, Parkinson’s disease, multiple sclerosis, and ischaemic stroke were obtained from UK Biobank PRS fields and z-standardized prior to analysis^37^(Supplementary Table 8).

Analyses were performed in participants with available PRS, organ level proteomic metrics, and covariate data (n = 40,867). For each disease PRS and each proteomic metric family, ordinary least squares regression models were fitted with standardized PRS as the outcome and organ level proteomic metrics as predictors. Models were adjusted for age, sex, and genetic ancestry using genetic principal components 1-10.

Two complementary model types were used. In multivariable PRS models, the same VIF-filtered predictor sets described above were used, with all retained organ level proteomics metrics within each metric family included simultaneously as predictors. In single metric PRS models, each organ level proteomics metric was tested separately. Predictors were standardized before modeling, so beta coefficients represent the association between a one standard deviation increase in the predictors and PRS, adjusted for age, sex, and genetic PC1-10.

For each model, beta coefficients, 95% confidence intervals, and p-values were extracted. Benjamini–Hochberg FDR correction was applied across all p-values in the multivariable PRS result table and separately across all p-values in the single metric PRS result table.

#### Bootstrap mediation analysis

Mediation analyses were performed using disease specific PRS as the exposure, organ level proteomic metrics as mediators, and incident disease as the outcome. Continuous PRS and organ metrics were standardized before analysis. Mediator models were fitted using linear regression, and disease outcome models were fitted using logistic regression. Models were adjusted for age, sex, and genetic PC1–PC10, with follow-up time additionally included in the outcome model. Indirect, direct, total effects, and mediated proportions were estimated using bootstrap resampling with 1,000 iterations.

### Computational Resources

All analyses were performed in Python v3.12 using Jupyter notebooks. Age gap, PCA, diagnosis, PRS, and phenotype association analyses were run in a local Python/Jupyter environment, whereas the PLS models were run on the UK Biobank Research Analysis Platform (UKB-RAP). Core Python libraries included NumPy, pandas, scikit-learn, statsmodels, matplotlib, and seaborn for data preprocessing, machine learning models, statistical analyses, and visualization. Recorded package versions included NumPy 2.1.0, pandas 2.2.3, scikit-learn 1.6.1, and matplotlib 3.9.2 in the local analysis environment.

## Results

### Organ level proteomic metrics capture overlapping but distinct biological signals

We first examined relationships among the three-organ level proteomic metrics: age gap, PC1, and PLS1. Pairwise correlations across organ-specific and global protein models showed a structured pattern of overlap among metrics, with correlations often observed between metrics derived from the same organ and between global protein models (Figure 2a). However, this overlap was not uniform across organs. Several organs showed clear cross-metric correspondence in the correlation matrix, whereas others, including esophagus, whole blood, and skin, showed weaker or more divergent relationships. These patterns indicate that the dominant proteomic axis captured by PC1 is variably related to age and organ-relevant physiology across organs. PLS1 metrics also showed relatively high correlations with each other and with certain PC1 metrics, consistent with the idea that organ-specific phenotypes covary with organ-specific proteomic signals. Comparisons within organ metrics also revealed insights. Global organ-specific protein age gap was most strongly correlated with brain and arterial age gap. This may reflect a link between aging of these systems and whole-body aging, though it is notable also that the brain contributed the largest proportion of proteins to the organ-specific protein signal (26.4% of total). Similarly, the dominant axis of variation (PC1) across all proteins was highly correlated with PC1 of immune, arterial and whole blood systems, highlighting the influence of these more distributed systems on whole body dynamics.

**Figure 2.**
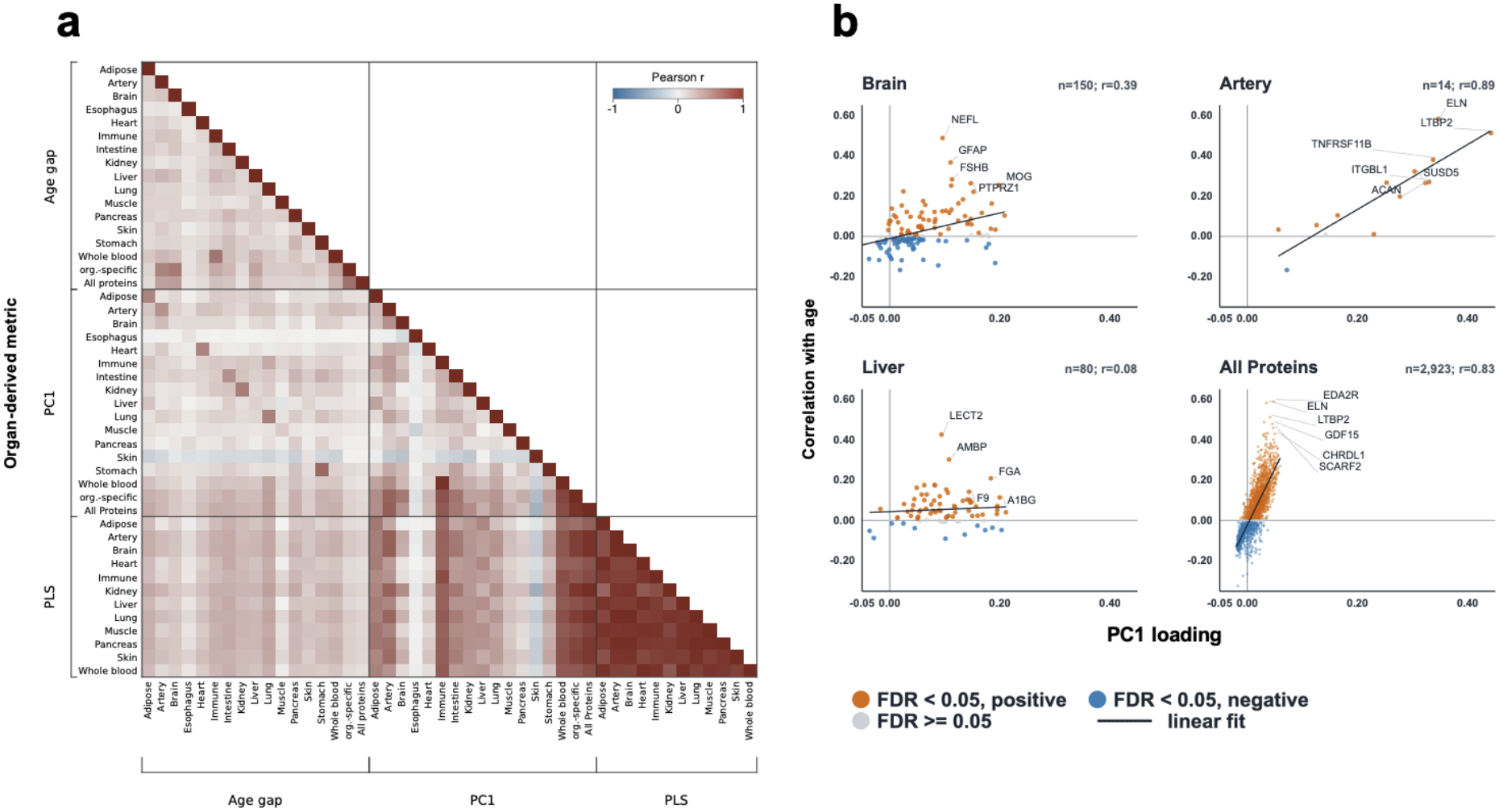
Relationships among organ-specific proteomic metrics. **a)** Correlation matrix of age gap, PC1, and PLS1 metrics across organs. Age gap represents predicted organ age relative to chronological age, PC1 represents the first principal component of organ-associated proteomic variation, and PLS1 represents a latent component derived from covariance between proteomic profiles and organ-relevant phenotypes. **b)** Scatterplots showing the relationship between PC1 loadings and protein–age associations for selected example organs, and for all proteins.

### Age is a major, but not always primary, effector of organ-specific proteomes

Organ age gap metrics are explicitly anchored to chronological age, but it is unclear whether age is a predominating signal in some or all organ proteomes. We therefore asked whether PCA-derived components reflected age-related protein variation, by correlating organ-specific PC1–PC3 loadings with protein age associations for each organ (Figure 2b; Supplementary Figure 2; Supplementary Table 7). After FDR correction, PC1 loadings were significantly aligned with protein age associations in 8 of 15 organ-specific, including artery, brain, immune, intestine, kidney, stomach, whole blood, and esophagus, although the esophagus model contained only three proteins and was interpreted cautiously, Global protein models also showed strong alignment between PC1 and age-related protein variation. In several organs where PC1 showed little or no age alignment, age-related variation instead emerged in higher order components, such that at least one of the first three PCs was significantly aligned with protein–age associations in 13 of 15 organ-specific models.

These findings indicate that age-related variation is a prominent feature of many organ-associated proteomes but is not necessarily their dominant source of variation. Thus, age anchored age gap metrics and unsupervised PCA-derived metrics capture related but distinct dimensions of proteomic variation.

### Proteomic organ metrics align with organ-relevant physiological phenotypes

To evaluate how organ level proteomic metrics relate to concurrent organ-specific physiological variation, we tested associations between age gap, PC1, and PLS1 metrics and organ-relevant phenotypes representing outward manifestations of organ structure or function (Figure 3a). Because PLS1 was derived using organ-relevant phenotypes, stronger phenotype associations for PLS1 were expected by design and were interpreted as phenotype guided alignment rather than unbiased superiority over other metrics (Supplementary Table 6).

**Figure 3.**
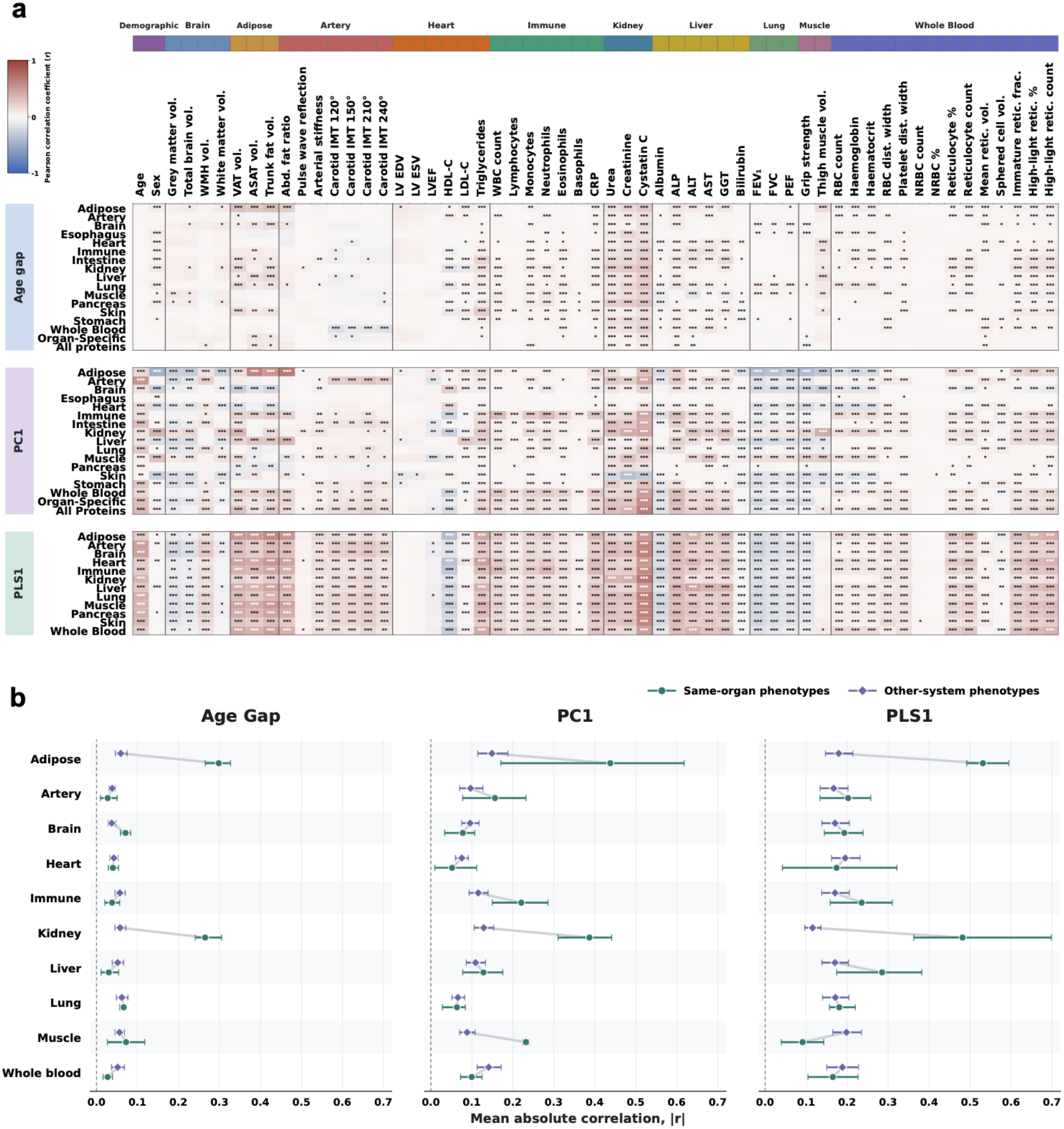
Phenotypic correlation landscape and organ-specificity of organ health metrics. **a)** Heatmap of Pearson correlation coefficients between organ health metrics; age gap, PC1, and PLS, and phenotypes. Rows represent metrics grouped by metric family and source organ, while columns represent phenotypes grouped by organ system. Asterisks indicate Benjamini–Hochberg-adjusted significance: *FDR < 0.05, **FDR < 0.01, and ***FDR < 0.001. **b)** Forest plots comparing mean absolute correlations (|r|) with phenotypes from the same organ system versus other systems across organs. horizontal bars show 95% bootstrap confidence intervals from 10,000 phenotype level resamples.

Across metric–phenotype pairs, median absolute correlations were highest for PLS1 (median |r| = 0.158), followed by PC1 (median |r| = 0.075) and age gap (median |r| = 0.036). Representative organ-relevant associations were observed across several systems. Adipose metrics were associated with adiposity measures, including adipose age gap with trunk fat volume (r = 0.327). Kidney metrics were strongly associated with renal markers, particularly kidney PLS1 (r = 0.698) and kidney age gap (r = 0.305) with cystatin C. Liver metrics were associated with liver enzymes and proteins, immune metrics with leukocyte traits, and brain PLS1 showed the strongest PLS1 association across all four brain related phenotypes. Several phenotypes were associated with metrics from multiple organs rather than exclusively with their assigned organ system. Cystatin C provided the clearest example: although kidney PLS1 showed the strongest association, several non-kidney PLS1 metrics also correlated strongly with cystatin C (r ∼0.62–0.65). This indicates that some physiological phenotypes reflect shared systemic proteomic variation in addition to organ matched signals.

We next compared associations with phenotypes assigned to the same organ system versus phenotypes assigned to other systems (Figure 3b). For all three metric classes, matched organ phenotypes had higher mean absolute correlations than other organ phenotypes; however, the scale and interpretation varied depending on the metric type. Age gap showed relatively modest associations overall, with mean |r| = 0.063 for matched organ phenotypes compared to 0.048 for other organ phenotypes. PC1 exhibited stronger effects, with mean |r| = 0.155 for matched organ and 0.095 for other organ phenotypes. PLS1 showed the highest overall correlations, with mean |r| = 0.230 for matched organ phenotypes and 0.173 for other-organ phenotypes. Thus, PLS1 showed the strongest absolute matched organ associations, but also substantial cross-organ associations, whereas PC1 showed a clearer relative separation between matched and other organ phenotypes. Matched organ enrichment was particularly evident for adipose, kidney, brain, liver, and immune phenotypes, although the degree of organ correspondence varied across systems and was not universal.

These results indicate that age gap showed weaker and more selective correspondence with concurrent physiology, PC1 captured broader physiological variation while retaining organ related structure, and PLS1 showed the strongest phenotype alignment but also substantial shared cross-organ variation. These findings suggest that age anchored, unsupervised, and phenotype guided proteomic metrics capture distinct dimensions of organ related biological variation, and that proteomic organ aging does not necessarily map directly onto concurrent organ-relevant physiological phenotypes.

### Organ level proteomic metrics show distinct associations with brain structure

We tested whether organ level proteomic metrics were associated with brain structural measures. We first tested each organ level proteomic metric separately for associations with cortical thickness and subcortical volume measures (Supplementary Figure 4), followed by multivariable models accounting for correlation among organ proteomics metrics.

The three proteomic metric families showed distinct patterns of association with brain structure (Figure 4). PC1 metrics showed several FDR-significant associations across cortical and subcortical measures, indicating that dominant unsupervised proteomic variation across several organ systems was independently related to brain structure. Among the PLS1 metrics, brain and immune PLS1 showed predominantly negative associations with cortical thickness, whereas skin PLS1 showed predominantly positive associations across several cortical regions. Brain age gap showed a more regionally restricted pattern of association than brain PLS1.

**Figure 4.**
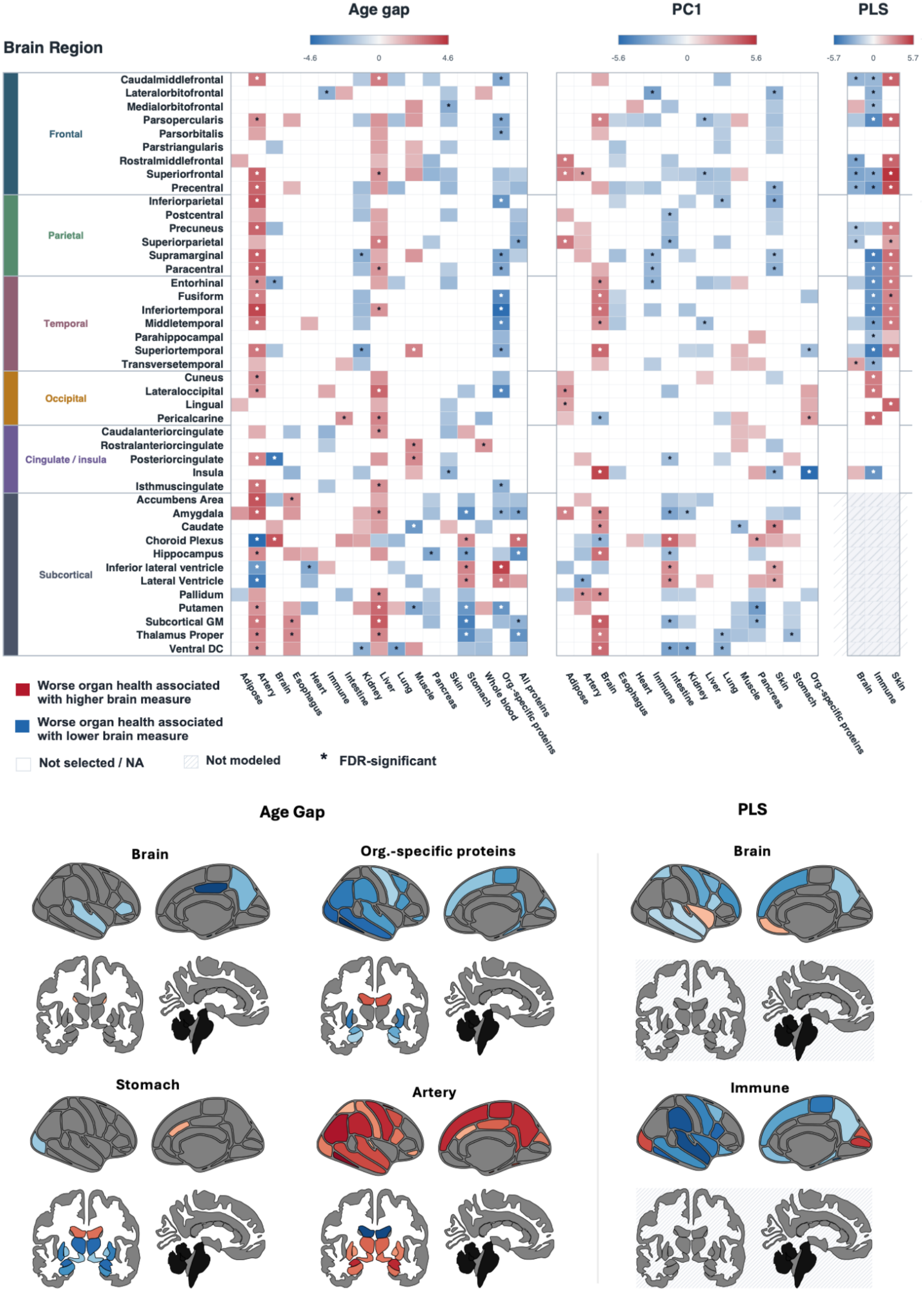
Associations between organ level proteomic metrics and brain structure. Heatmaps showing retained associations between organ level proteomic metrics and cortical thickness or subcortical volume measures from multivariable regression models adjusted for age, sex, and imaging center, with eTIV additionally included for subcortical volume models. Columns are grouped by proteomic metric type: age gap, PC1, and PLS1. For PLS1, only brain, immune, and skin metrics are shown because multicollinearity among PLS1 organ scores led to the use of a reduced candidate set in multivariable models. Rows represent cortical and subcortical brain measures grouped by anatomical region. Colors indicate the direction and magnitude of model statistics for retained associations, and asterisks indicate FDR-significant associations. Blank cells indicate metrics that were not selected or were not available; hatched cells indicate measures that were not modeled. Brain maps present selected examples of spatial association patterns for brain-derived and non-CNS organ level proteomic metrics.

Several non-CNS organ age gap metrics showed FDR-corrected associations with brain structural measures, indicating that systemic organ-specific proteomic aging is related to brain structure beyond brain-derived metrics alone. Higher muscle age gap was independently associated with lower volumes of the caudate and putamen, which is significant considering the involvement of these structures in motor circuits. Greater stomach age gap was specifically associated with smaller volumes in subcortical and medial temporal lobe regions and enlarged ventricles, but not with cortical areas, indicating a pattern more pertinent to subcortical structures. The all organ-specific protein age gap metric was also independently associated with widespread cortical thinning and larger ventricular volumes, suggesting a broader systemic proteomic aging component in addition to individual organ effects.

Unexpected positive associations were also observed for selected organ level proteomic metrics. These associations suggested that organs with more advanced age or worse function were paradoxically associated with thicker cortex or larger brain volumes. This phenomenon was most prominent and consistent with artery and liver age gap, and with immune and skin PLS1 model. While these findings are challenging to interpret, closer examination suggests multiple explanations. In univariate models (Supplementary figure 4), immune and skin PLS1 metrics show the expected negative relationship with cortical thickness. Given the high collinearity between PLS metrics (Fig. 2A), the positive relationships observed in the multivariate models likely represent small but significant residual effects, or perhaps complex protein-protein interactions. In contrast, artery and liver age-gaps show positive relationships with brain thickness/volume even in univariate analyses. Given the well documented relationship between e.g. vascular aging and brain integrity, these paradoxical positive relationships may represent survivor effects. The artery age gap model is one of the most reliable models, so this result cannot be attributable to model quality (Supplementary table 4). However, these estimates should be interpreted cautiously and considered exploratory until replicated.

### Age anchored proteomic metrics show stronger associations with future neurodegenerative diagnoses

We next tested whether baseline organ proteomic metrics were associated with subsequent ICD-10 defined clinical diagnoses using single metric and multivariable Cox proportional hazards models (Figure 5a; Supplementary Figure 5)

**Figure 5.**
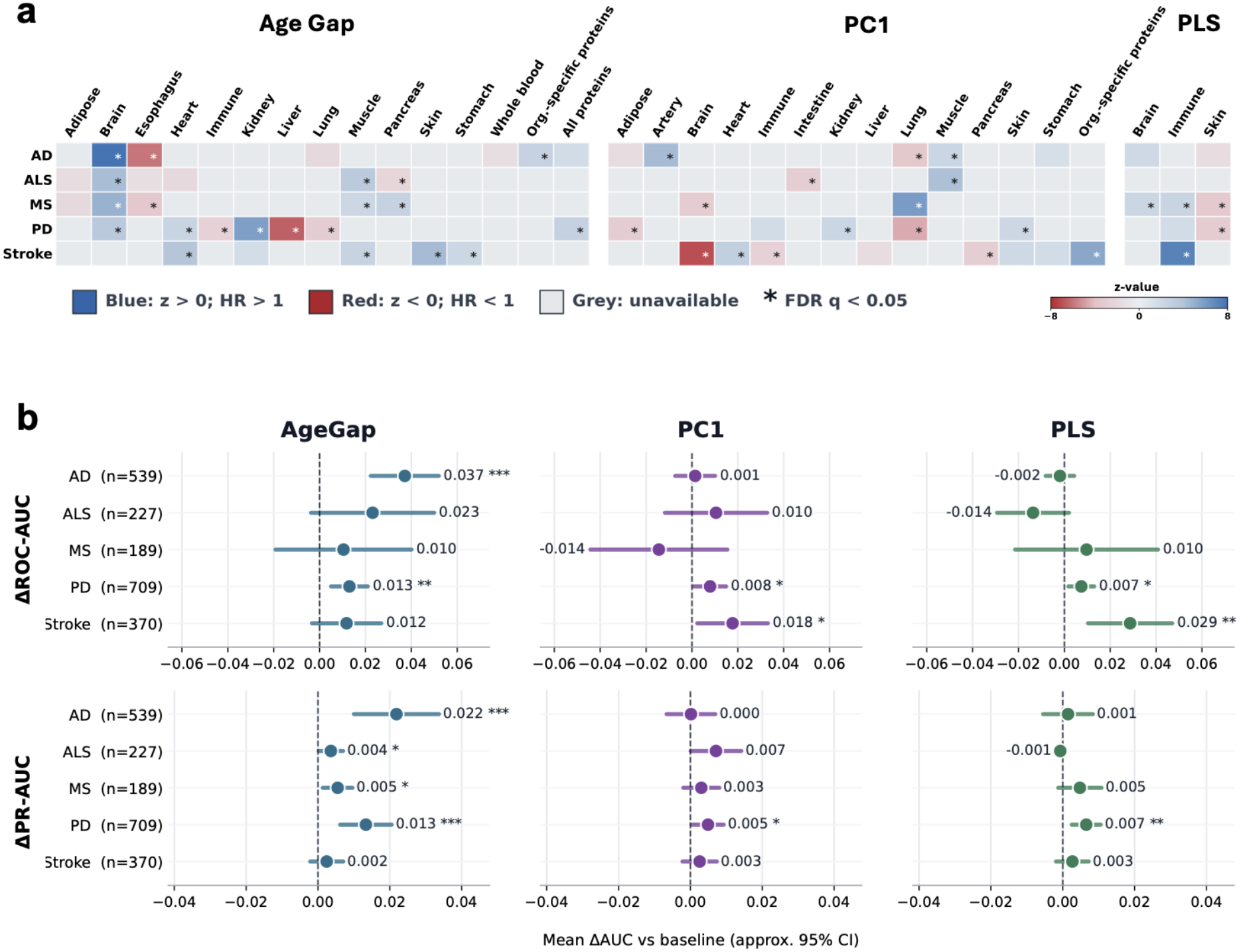
Associations and incremental predictive value of organ level proteomic metrics for later neurological diagnoses. **a)** Heatmaps show associations between organ level proteomic metrics and later ICD-10-based diagnoses of Alzheimer’s disease (AD), amyotrophic lateral sclerosis (ALS), multiple sclerosis (MS), Parkinson’s disease (PD), and stroke using Cox proportional hazards models. Results are shown separately for age gap, PC1, and PLS1 metrics. Colors represent Cox Wald z-statistics, with blue indicating positive associations/higher hazard and red indicating negative associations/lower hazard. * indicate FDR-significant associations (PFDR < 0.05). Models were adjusted for age and sex, with follow-up time used as the Cox time-to-event variable. **b)** Incremental predictive value of organ level proteomic metrics for later neurological diagnoses. Regularized logistic regression models were evaluated using repeated stratified cross-validation. For each outcome, a baseline model including age, sex, and follow-up time was compared with expanded models additionally including all organ age gap, PC1, or PLS1 metrics. Points show mean ΔAUC, defined as expanded model AUC minus baseline model AUC, and horizontal bars show approximate 95% confidence intervals based on corrected resampled standard errors from fold-level ΔAUC values. The top row shows ΔROC-AUC and the bottom row shows ΔPR-AUC, where PR-AUC denotes precision recall AUC. Positive values indicate improved discrimination after adding organ level proteomic metrics. Asterisks indicate corrected resampled t-test p-values.

Age gap, PC1 and PLS1 metrics showed different association patterns with later clinical diagnoses. Models using a single metric showed broader disease association patterns across organ systems, whereas multivariable models retained a smaller set of organ metrics, consistent with shared variance among organ level proteomic signals. Age gap metrics showed the clearest disease related patterns, including associations of brain age gap with incident Alzheimer’s disease and ALS, muscle age gap with incident ALS, heart age gap with incident stroke, and heart and kidney age gap with incident Parkinson’s disease. In contrast, PC1 and PLS1 measures exhibited sparse and more heterogeneous associations with diagnostic outcomes.

Next, we compared baseline models that included age, sex, and follow-up time with repeated cross-validated models that included each proteomic metric family to see if these disease connections translated into incremental predictive value (Figure 5b). The overall improvement in prediction was modest, even if there were a few statistically significant increases. The biggest gains were seen in the age gap models, especially for Parkinson’s and Alzheimer’s diseases, but the associated rises in ROC-AUC and PR-AUC were still quite small. The enhancements across diagnoses were more variable and smaller for the PC1 and PLS1 models. These findings collectively show that age anchored organ proteomic metrics show stronger and more consistent associations with subsequent neurodegenerative and vascular diagnoses than unsupervised or phenotype guided summaries, but their limited incremental predictive performance implies that they are not sufficient as stand-alone biomarkers for neurodegenerative diagnosis in the future.

To provide additional context to these disease associations, we tested whether organ level proteomic metrics were associated with polygenic risk scores for Alzheimer’s disease, Parkinson’s disease, multiple sclerosis and stroke (Figure 6a, Supplementary Figures 6 and 7). PRS associations were generally small but displayed several disease specific patterns. Alzheimer’s disease PRS was associated with brain and immune PC1, brain PLS1, and global organ-specific age gap metrics, whereas stroke PRS exhibited the broadest pattern of associations across age gap, PC1, and PLS1 metrics. The direction of association differed across brain derived metrics, consistent with these measures capturing non-equivalent dimensions of proteomic variation. The Parkinson’s disease PRS was most associated with PC1 and PLS1 measures such as PC1 for liver and skin and PLS1 for brain, immune system and skin. PRS for multiple sclerosis was associated with specific age gap and PC1 measures including immune and heart age gap, skin age gap and liver and skin PC1. Taken together, the PRS analyses indicate that organ level proteomic metrics capture disease relevant variation that is partially overlapping with genetic liability but is also metric and disease specific. These PRS results imply some overlap of organ level proteomic metrics with inherited disease liability, but the patterns were metric and disease specific and should be interpreted as complementary genetic context, not validation of diagnostic prediction.

**Figure 6.**
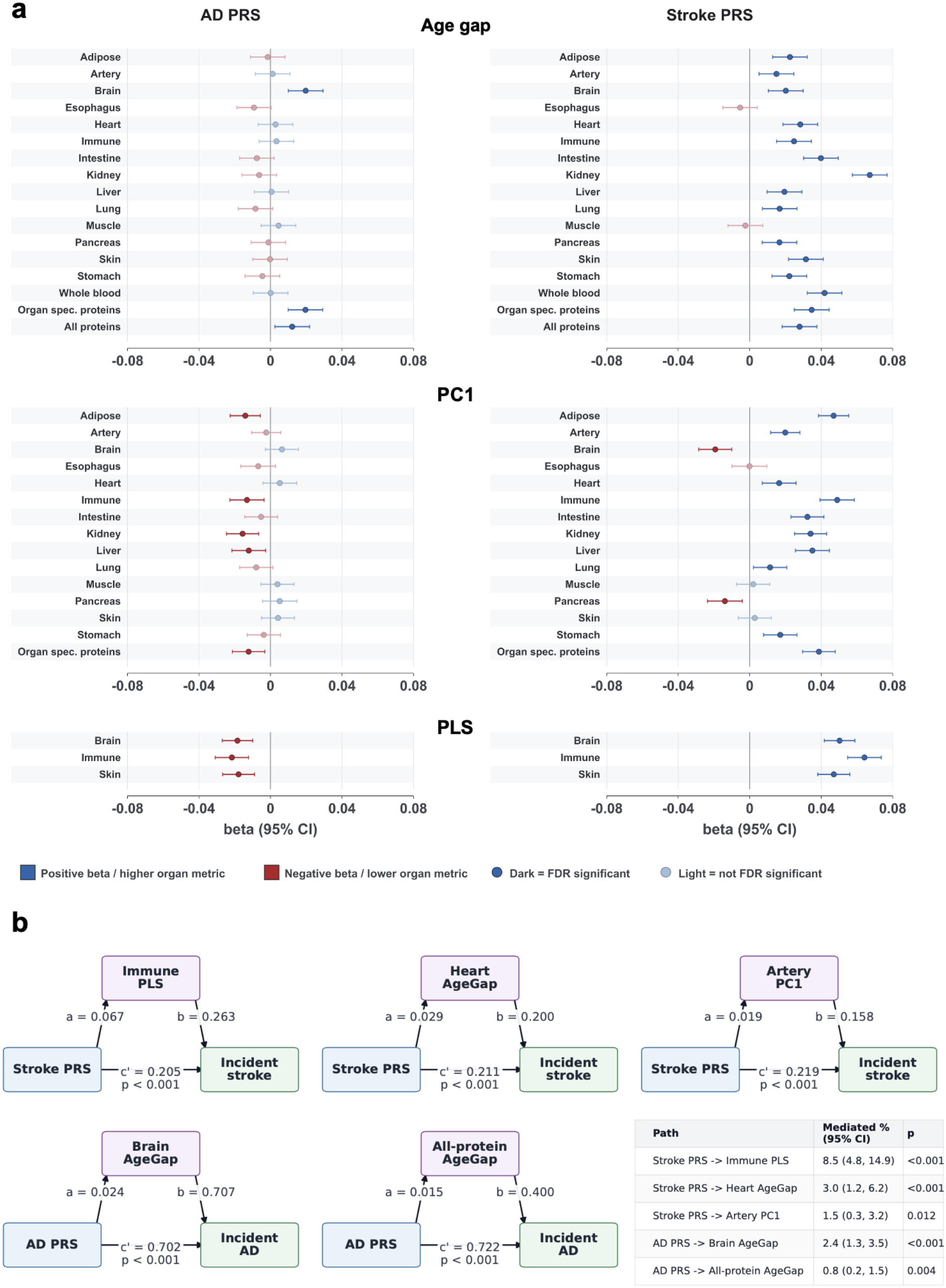
Genetic associations and mediation analyses linking organ level proteomic metrics to neurological disease risk. **a)** Forest plots show associations between disease polygenic risk scores (PRS) and organ level proteomic metrics from multivariable linear models. Results are shown for Alzheimer’s disease (AD) and stroke PRS across age gap, PC1, and PLS1 metrics. Points show standardized beta estimates, and horizontal bars indicate 95% confidence intervals. Blue indicates positive associations with higher metric values, while red indicates negative associations with lower metric values. Darker pointranges indicate FDR-significant associations. **c)** Path diagrams illustrate significant statistical mediation pathways linking disease PRS to incident AD or stroke through selected organ level proteomic metrics. Each panel shows the exposure PRS, mediator, and incident disease outcome, with path coefficients shown for the PRS-to-mediator path, mediator-to-outcome path, and direct PRS-to-outcome path. The table summarizes the percentage of the total PRS association mediated by each organ metric, with 95% bootstrap confidence intervals and mediation p-values. Mediator models were adjusted for age, sex, and genetic principal components 1–10; outcome models were additionally adjusted for follow-up time.

As a final pathway-oriented analysis, we tested whether selected organ level proteomic metrics statistically mediated associations between disease specific PRS and incident disease (Figure 6b). Bootstrap mediation analyses identified significant but modest indirect effects for selected pathways, including stroke PRS through immune PLS1, heart age gap, and artery PC1 to incident stroke, and AD PRS through brain age gap and all-protein age gap to incident Alzheimer’s disease. The strongest mediated proportion was observed for stroke PRS through immune PLS1 to incident stroke (8.5% [4.8%-14.9%]), although mediated proportions were modest across all tested pathways. These results suggest that organ level proteomic metrics capture small but detectable components of genetic liability–disease associations, supporting their interpretation as partial systemic intermediates rather than dominant explanatory pathways.

## Discussion

Biological age measures are increasingly used to estimate how biological systems deviate from chronological aging^15,38^. A related but distinct goal is to quantify variation in the molecular state of specific organ systems, which may reflect age-related change, organ function, systemic physiology, or other sources of biological variation^2,3,39^. Our findings show that these sources of variation are not easily separated in plasma proteomic data. Although organ age gap models provide an intuitive way to measure age-related deviation, they capture only one dimension of organ level proteomic biology^1^. By comparing age gap with unsupervised PC1 and phenotype guided PLS1 metrics across 15 organ systems, we found that the biological interpretation of an organ level metric depends strongly on how it is constructed. Global and organ-specific protein models showed a broad age-related proteomic signal, but organ-specific PC analyses indicated that this signal was not always the dominant axis of variation within individual organ protein sets. Phenotype-guided PLS1 metrics were more closely linked to organ-relevant physiological traits, whereas age gap metrics showed clearer associations with later clinical diagnoses. All three metrics families were associated with brain structure but showed distinct spatial and organ-specific patterns. Incremental predictive value for future diagnosis, and mediation of genetic risk was modest. Thus, although age gap metrics may be useful research tools for studying age-related organ–brain relationships, their limited predictive performance and diagnostic specificity do not support their use as stand-alone disease biomarkers. More broadly, our findings point to strong links between CNS and non-CNS proteomic aging signals, suggesting that brain health is embedded within systemic biological processes rather than being fully separable from peripheral organ biology.

Previous multi-organ proteomic aging studies have shown that organ age gaps vary across individuals and are associated with disease risk, mortality, health span, and longevity^1–3^. A key interpretive challenge, however, is that organ-specific age gaps may not always reflect pure organ-intrinsic biology. Since many circulating proteins change with age across multiple physiological systems, an organ-specific clock may capture part of a shared systemic aging signal, especially when organ models are analyzed in isolation. Our results support this concern; we found strong age-related structure in global all-protein and organ-specific proteomic metrics and found several organ level metrics to be correlated with these broader signals. This points towards the possibility that tissue-specific aging associations are driven in part by shared proteomic aging variance rather than independent tissue effects. Future studies should aim to isolate these overlapping signals and, as much as possible, analyze them independently.

A key aspect of this study is the structured comparison of three complementary organ level proteomic metrics. While previous organ proteomic aging studies have largely focused on age-gap models, we introduced unsupervised PC1 and phenotype-guided PLS1 as additional ways to summarize the organ-associated plasma proteome and compared these approaches directly with the more traditional age-gap framework. Our results indicate that, instead of being interchangeable markers of biological aging, all three metrics capture overlapping but distinct dimensions of proteomic variation. Age gap ties proteomic variation to chronological age, PC1 describes the first dimension of variation among organ-associated proteins, and PLS1 summarizes variation associated with organ-relevant phenotypes. This distinction matters because not all biological organ variation is necessarily age-related, and age-related variation is not always the dominant proteomic axis. Therefore, the choice of metric should depend on the biological question: age-related disease association, dominant molecular variation, or phenotype-linked organ biology.

In our study, age gap metrics showed more consistent associations with subsequent clinical diagnoses than PC1 or PLS1, suggesting that age-anchored proteomic variation contains disease-relevant information. This may also relate to the fact that the single biggest risk factor for most neurodegenerative diseases is age, and this work supports the idea of molecular aging playing some role in disease risk. However, the interpretation of age gap is shaped by its training objective; because these models are optimized to predict chronological age, they preferentially capture proteomic variation that covariates with age. This is useful when disease-relevant biology is age-related, as is the case with neurodegenerative diseases, but it may capture less of the organ-relevant variation that is only weakly related to chronological age. This may explain why age gap metrics demonstrated stronger disease associations, whereas PC1 and PLS1 showed stronger or more selective relationships with physiological phenotypes^3^. This distinction may provide a useful tool for researchers to discern which aspects of organ biology are canonical features of human aging vs. aspects that are driven by person-specific genetic risk and environmental exposures. Although the association between baseline age-anchored proteomic variation and later diagnosis is biologically compelling, the limited incremental predictive value of these models suggests that organ age gap metrics are unlikely to function as stand-alone biomarkers for neurodegenerative disease. Rather, they may be informative research tools for characterizing disease-associated age-related biology, without establishing diagnostic utility or disease mechanism.

Unsupervised PCA helped distinguish dominant proteomic variation from age-related variation. Previous work shows that aging is accompanied by coordinated multi-omic and systemic molecular changes, which can appear as shared axes of variation across biological datasets^40–42^. PC1 loadings were highly correlated with protein-age associations in global all-protein and organismal protein models, suggesting that a large fraction of the variation in the circulating proteome reflects an age-related systemic signal. But this pattern was not the same for all organs: some organ-specific protein sets were strongly age-aligned in PC1, while in others, age-related variation appeared in PC2 or PC3 was weaker overall. Adipose and heart showed the weakest age-related structure, with none of the first three PCs significantly aligned with protein-age associations after FDR correction. This is consistent with the largest published organ-specific plasma proteomic aging resource to date, in which the adipose and heart panels were built from the fewest organ-enriched circulating proteins of the 11 organs modelled (five and ten proteins, respectively)^1^, supporting our interpretation that limited protein coverage may partly explain the weaker age-related structure observed here. Beyond coverage, adipose- and heart-derived circulating proteins may also be disproportionately shaped by metabolic and cardiovascular physiology rather than chronological age per se. Obesity and cardiovascular ageing share overlapping molecular hallmarks, including senescence, mitochondrial dysfunction, and inflammation, that are not strictly age-dependent^43^, and multi-organ ageing atlases show that immune and metabolic remodeling in adipose tissue can be driven by factors other than chronological age^44^. In contrast, liver, lung, muscle, pancreas, and skin showed age-related alignment in secondary components rather than PC1, suggesting that age-related proteomic variation was present but not the dominant axis of variation. This pattern is broadly consistent with reports that non-chronological-age exposures can dominate the circulating proteomic signal in specific organs. Lung aging, for example, is strongly and specifically linked to smoking exposure in plasma proteomic and genetic studies^45^, and tissue-resolved transcriptomic surveys across the human lifespan likewise show that the magnitude and dominant drivers of age-related change differ substantially across liver, muscle, and other peripheral organs^46^. For these organs, PC1 may capture overlapping age-related, physiological, metabolic, inflammatory, and other sources of proteomic variation, rather than representing a purely age independent biological axis. Overall, chronological age represents an important source of circulating proteomic variation, but its prominence differs across organ-associated protein sets and may overlap with other physiological dimensions. At the same time, correlations between organ level metrics and global proteomic signals indicate that organ-associated patterns may contain shared systemic variation rather than fully independent organ-specific biology. Distinguishing between dominant organ level variation, age-related variation, and shared systemic proteomic aging signals is therefore important for interpreting associations between organ level proteomic metrics, brain structure, and disease.

Systemic organ biology provides an important context for interpreting associations between plasma proteomic metrics and brain structure. Prior work has linked cardiovascular, renal, metabolic, and immune health to brain aging and neurodegenerative vulnerability^14^, supporting the idea that brain health is shaped by both CNS and non-CNS processes. These links may arise through several partially overlapping pathways, including vascular dysfunction, altered cerebral perfusion, systemic inflammation, impaired metabolic homeostasis, reduced protein clearance, and changes in blood-brain barrier integrity^14^. In our analyses, non-CNS organ level proteomic metrics remained in multivariable models for several cortical and subcortical brain measures, suggesting that systemic proteomic variation is related to specific brain structures beyond metrics derived from the brain alone. These findings align with evidence demonstrating widespread associations between individual plasma proteins and brain imaging measures in the UK Biobank, highlighting links with neurobiological and immune related processes^47^. However, interestingly, we find that proteins from certain organs are associated with specific brain structures. Some of these associations highlighted known brain-body links. Older muscle proteomic age was associated with lower caudate and putamen volumes consistent with the established role of the striatum in learning and execution of motor movement^48^. The relationship between skin proteomic PC1 and somatomotor cortex was interesting given that this network includes the primary somatosensory cortex, which receives signals originating in peripheral nerves with terminals in the skin^49^. The association between older pancreas age and the hippocampus was an interesting observation given the central role of the pancreas in glucose regulation and frequent reports of diabetes being a risk factor for Alzheimer’s disease^50^. However, some relationships warrant further investigation such as the relationship between the stomach and subcortical volumes, supported by both age gap PC1. These results suggest the presence of multiple brain-body axes that age at different rates.

In disease analyses, heart and kidney age gap metrics persisted within models of later Parkinson’s disease, consistent with epidemiological studies associating heart failure and chronic kidney disease to increased risk of Parkinson’s disease^51,52^. This is corroborated by direct evidence of organ-specific synuclein pathology: cardiac sympathetic denervation and postmortem cardiac α-synuclein deposition are well documented in Lewy body disease^53,54^, These findings may reflect shared vascular, inflammatory, metabolic, or clearance-related pathways rather than direct organ-to-brain causality. For example, cardiovascular dysfunction could influence brain vulnerability through perfusion and vascular injury, whereas kidney dysfunction may reflect impaired clearance, systemic inflammation, or broader cardiometabolic burden^13,55^. Peripheral protein handling and peripheral proteinopathy have also been suggested as mechanisms linking systemic biology and neurodegenerative disease, although our plasma proteomic analyses cannot directly test these mechanisms. Collectively, these findings are consistent with a systems-level model where multiple peripheral processes may be related to brain vulnerability but underscore that the biological interpretation is highly dependent on how proteomic variation at the organ level is summarized.

More broadly, these results raise the question of what “proteomic aging” measured in plasma represents. Plasma proteins are not a direct measurement of isolated organ tissue state; they reflect secretion, leakage, immune activation, vascular and metabolic physiology, tissue remodeling, protein clearance, and contributions from multiple organs^22,56^. Therefore, plasma proteomic aging is unlikely to represent only a slow accumulation of inflammatory or degradative damage. Instead, it may reflect a composite systemic state in which broad aging-related processes coexist with more specific organ-derived and phenotype-linked signals^15^. This interpretation is supported by our observation that global proteomic aging related to brain structure, Alzheimer’s disease, and stroke genetic liability, while individual organ metrics captured more selective associations. Thus, plasma proteomic aging may be best viewed as a distributed systems-level readout of biological state rather than a direct measure of isolated organ deterioration.

The major strengths of this study include its large-scale cohort design, integration of multiple data modalities, multivariate design, and direct comparison of alternative organ level proteomic metrics. Several limitations should be noted in the interpretation of these findings. First, the analyses were observational and cannot establish causal ordering between organ level proteomic variation, brain structure, and subsequent clinical diagnosis. The diagnostic analyses used follow-up data, but should be considered as longitudinal associations, not validated prediction models or disease mechanisms. In addition, plasma proteomics and brain MRI were acquired at different UK Biobank assessment visits and were therefore not contemporaneous. As a result, the proteomic and imaging measures may represent different biological time points, introducing temporal mismatch into the observed proteomic brain structure associations. Second, participants of the UK Biobank are healthier than the general population, which may limit generalizability and may attenuate associations in more clinically diverse populations^57^. Third, plasma proteomics provides an indirect readout of organ biology. Circulating protein levels may reflect secretion, clearance, inflammation, transport, or contributions from multiple tissues, rather than molecular activity within a single organ^56^. Relatedly, organ-specific protein mapping was based on tissue-enriched gene expression annotations, and gene expression does not necessarily correspond to circulating protein abundance^58^. This strategy also excludes many proteins expressed across multiple organs, which may be biologically important for systemic aging and inter-organ communication^59^. The phenotype guided PLS1 metrics were also highly correlated across organ systems, indicating that they capture substantial shared proteomic variation in addition to organ-related signals. This limits the interpretation of individual PLS1 coefficients as fully organ-specific effects, particularly in multivariable models. Fourth, some organ panels were relatively small in size, limiting stability of organ-specific estimates. Finally, case numbers were limited for some neurological diagnoses, particularly rarer disorders, and all analyses were conducted in a single cohort. Independent datasets with similar proteomic, imaging and longitudinal clinical data will be required to assess generalizability by replication.

Future work should test these organ level proteomic metrics in longitudinal cohorts with repeated proteomic, imaging, cognitive, and clinical measures. Such studies would help to determine whether age-anchored, unsupervised, and phenotype-guided metrics capture stable biological traits, dynamic change over time, or early signals associated with later clinical decline. Replication in more diverse populations will also be important for evaluating generalizability across genetic, environmental, ethnic and socioeconomic contexts. Finally, the integration of organ level proteomic metrics into interventional or mechanistic studies may help clarify whether systemic pathways such as inflammation, metabolism, vascular function, or protein handling contribute to brain-related outcomes.

In summary, we show that plasma proteomic organ metrics do not reflect a single biological construct. Age gap, PC1 and PLS1 are related but distinct axes of organ level proteomic variation, exhibiting differential associations with chronological age, physiological phenotypes, brain structure and subsequent clinical diagnoses. These findings support that brain related outcomes are associated with proteomic variation across both CNS and non-CNS systems, supporting a distributed brain body perspective on neurodegenerative disease. At the same time, they show that the biological interpretation of organ level proteomic metrics depends strongly on how those metrics are defined.

## Data availability Statement

All data in this manuscript was sourced from the UKBiobank, under Project ID number 105777.

## Code availability statement

Code used for the main analyses presented in this study can be found at https://github.com/DeMONLab-BioFINDER/Zendehdel_Organ_Health.git.

## Author contributions

RZ and JWV conceptualized the study. RZ, LA, LC, GV, and JWV contributed to analytical design. RZ, LA, JS and RB were involved in data processing and curation. RZ conducted data analysis. JWV supervised the study. RZ and JWV wrote the manuscript. All authors interpreted the data and contributed substantively to revising the manuscript.

## Acknowledgements

This work was funded by the SciLifeLab & Wallenberg Data Driven Life Science Program (grant: KAW 2020.0239) and the Swedish Research Council (2024-03642). This research has been conducted using the UK Biobank Resource under application number 105777.

