## Supplementary Information for "Beyond organ age gaps: comparing plasma proteomic metrics of organ biology in relation to brain structure and neurodegenerative disease"

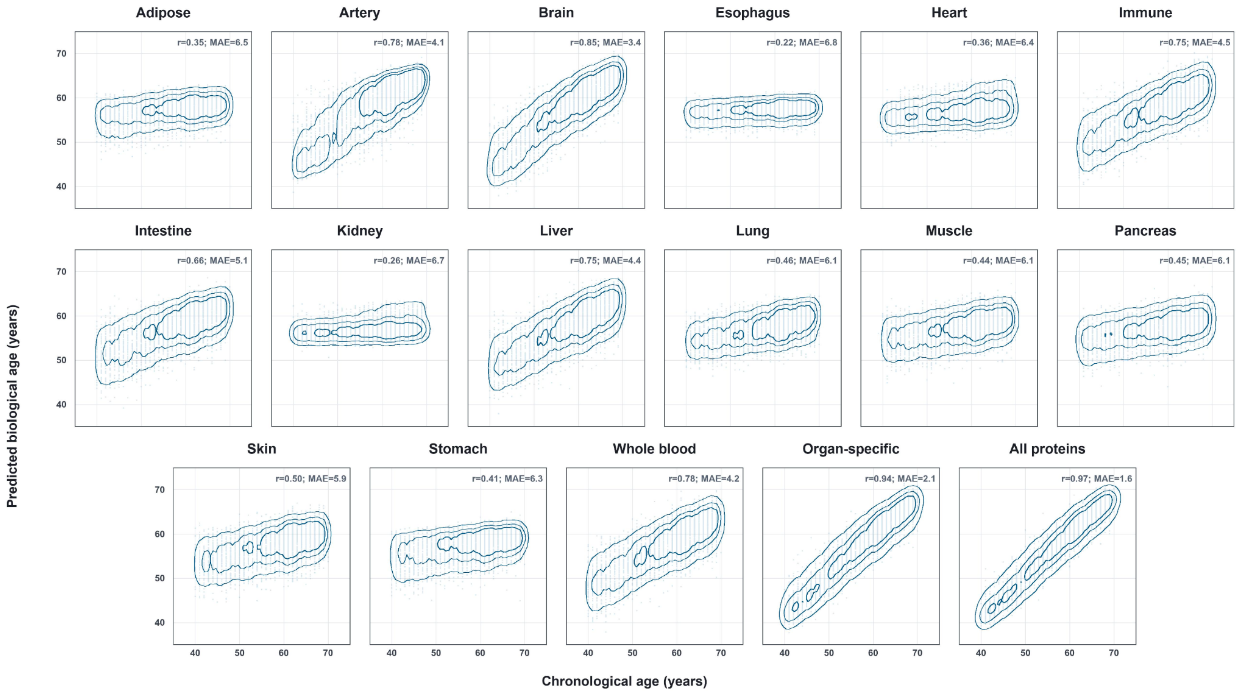


**Supplementary Figure 1. Model calibration of organ-specific age prediction models.** Predicted biological age is plotted against chronological age for each organ-specific XGBoost model in the independent test set. Density contours illustrate the distribution of predictions across the age range, enabling visual assessment of model calibration, age-dependent bias, and overall agreement between predicted and observed age prior to age-gap residualization.

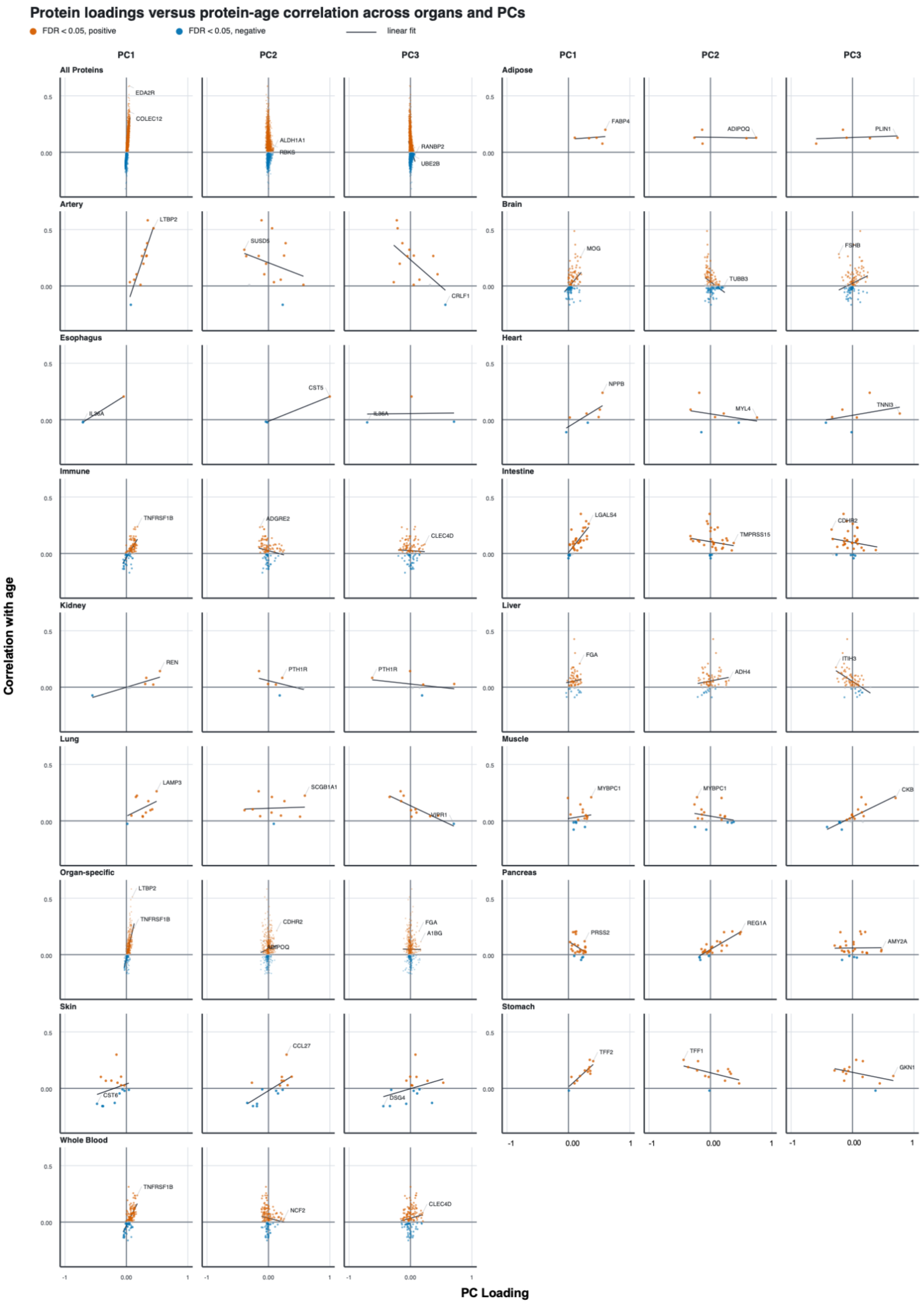
Protein loadings versus protein-age correlation across organs and PCs
FDR < 0.05, positive
FDR < 0.05, negative
linear fit
PC1
PC2
PC3
All Proteins
Adipose
Artery
Brain
Esophagus
Heart
Immune
Intestine
Kidney
Liver
Lung
Muscle
Organ-specific
Pancreas
Skin
Stomach
Whole Blood
EDA2R
COLEC12
ALDH1A1
RBKS
RANBP2
UBE2B
FABP4
ADIPOQ
PLIN1
LTBP2
SUSD5
CRLF1
MOG
TUBB3
FSHB
IL36A
CST5
NPPB
MYL4
TNNI3
TNFRSF1B
ADGRE2
CLEC4D
LGALS4
TMPRSS15
CDHR2
REN
PTH1R
FGA
ADH4
ITIH3
LAMP3
SCGB1A1
VIPR1
MYBPC1
CKB
CDHR2
ADIPOQ
A1BG
PRSS2
REG1A
AMY2A
CST6
CCL27
DSG4
TFF2
TFF1
GKN1
NCF2
Correlation with age
PC Loading
0.5
0.00
-1
1

**Supplementary Figure 2. Organs correlations between PCA components and age-related protein variation.** Each panel shows protein-level scatterplots for PC1, PC2, and PC3 across 15 organ systems and global protein sets. The x-axis represents the correlation of each protein with age, and the y-axis represents the corresponding PCA loading. PC1 demonstrates a consistent and statistically significant alignment with age-associated proteomic variation in multiple organs, whereas PC2 and PC3 generally exhibit weaker or inconsistent associations. Regression slopes and p-values reflect the strength of association within each organ.

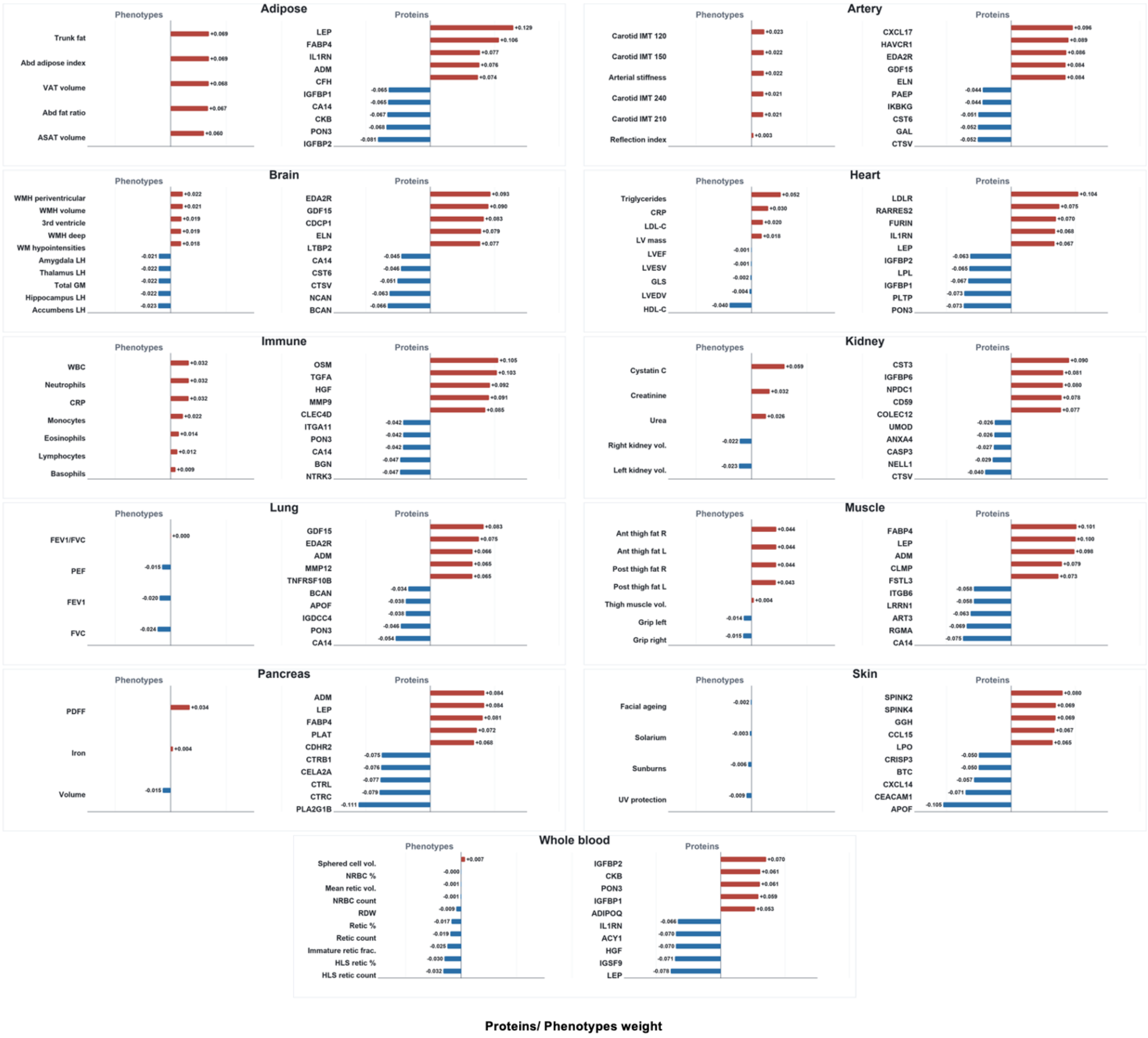


**Supplementary Figure 3. PLS feature weights by organ.** Top positive and negative PLS1 feature weights are shown for each organ-specific PLS model. Within each organ panel, phenotype weights and protein weights are displayed side by side using a shared x-axis scale to enable comparison of loading magnitudes across organs and feature classes. Red bars indicate positive PLS1 weights and blue bars indicate negative PLS1 weights.

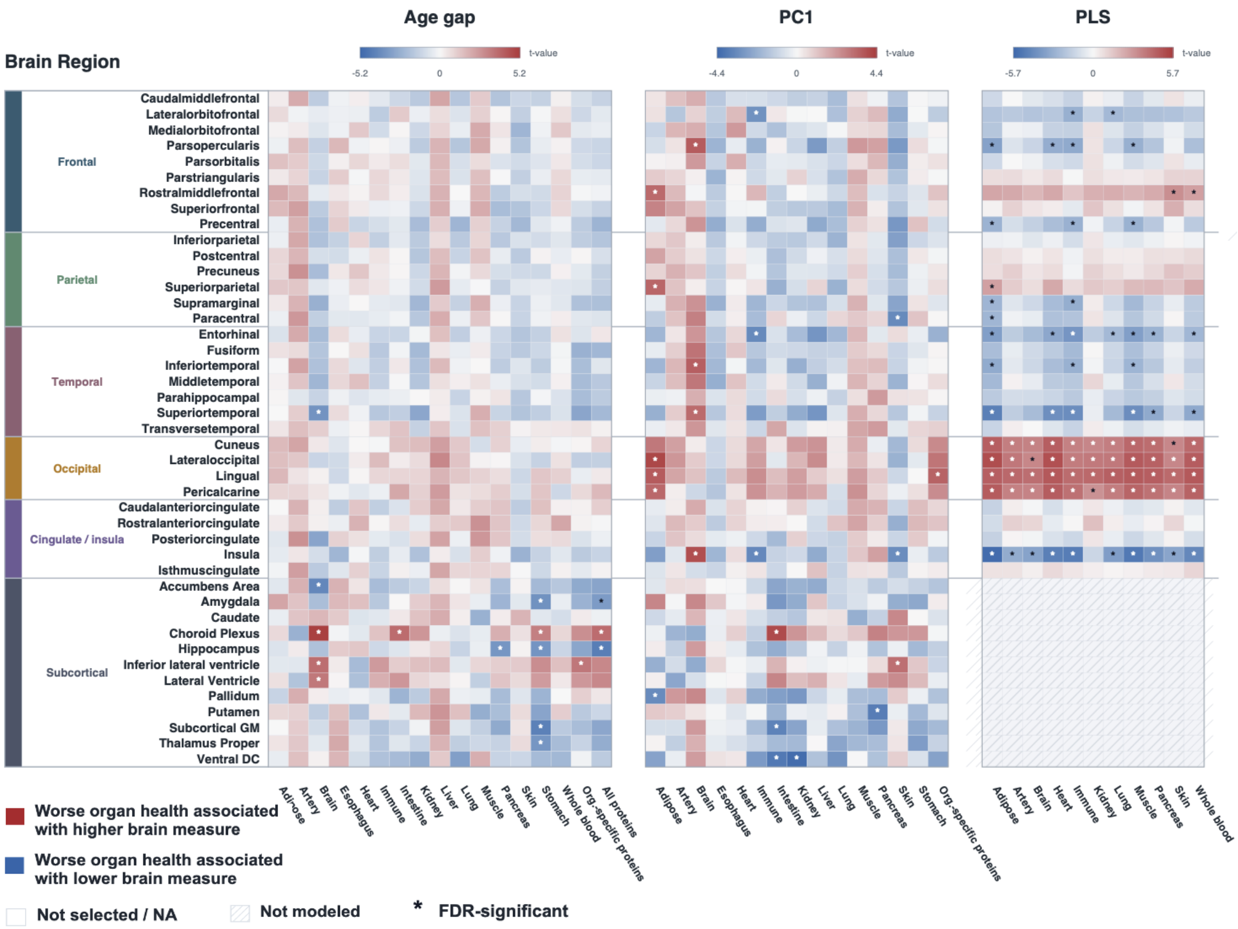


**Supplementary Figure 4. Single-metric associations between organ level proteomic metrics and brain structure.** Heatmaps show associations between individual organ level proteomic metrics and regional cortical thickness or subcortical volume measures. Models were fitted separately for each organ metric and brain structural outcome. Columns are grouped by metric type: age gap, PC1, and PLS1. Rows represent cortical and subcortical brain measures grouped by anatomical region. Colors indicate signed model t-values, with red indicating higher metric values associated with higher brain measures and blue indicating higher metric values associated with lower brain measures. Asterisks indicate FDR-significant associations. Blank cells indicate missing or unavailable estimates, and hatched cells indicate outcomes that were not modeled. PLS1 subcortical associations were not modeled.

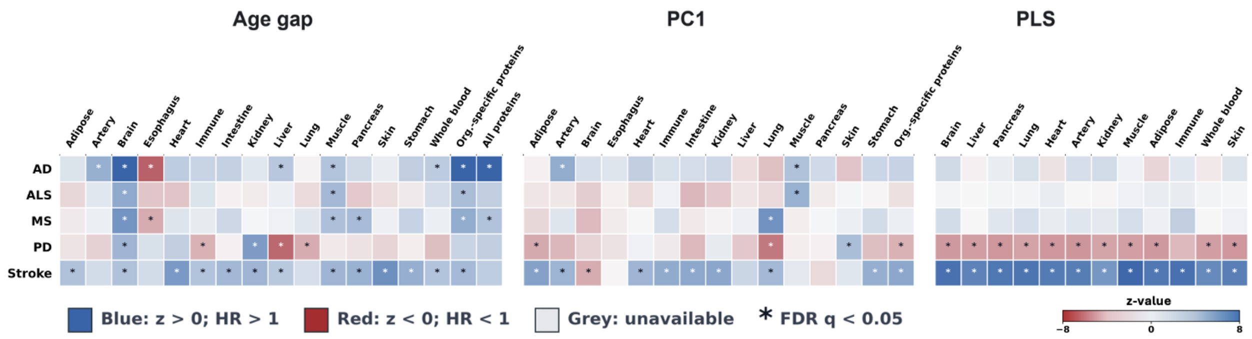


**Supplementary Figure 5. Single-metric associations between organ level proteomic metrics and later clinical diagnoses.** Heatmaps show associations between individual organ level proteomic metrics and later ICD-10-based diagnoses of Alzheimer's disease, amyotrophic lateral sclerosis, multiple sclerosis, Parkinson's disease, and stroke. Cox proportional hazards models were fitted separately for each organ metric and diagnostic outcome, adjusting for age and sex. Follow-up time was used as the Cox time-to-event variable and was not included as a covariate. Columns are grouped by metric type: age gap, PC1, and PLS1. Colors indicate Cox Wald z-statistics, with blue indicating positive associations with higher disease hazard (HR > 1) and red indicating negative associations with lower disease hazard (HR < 1). Asterisks indicate FDR-significant associations. Grey cells indicate missing or unavailable estimates.

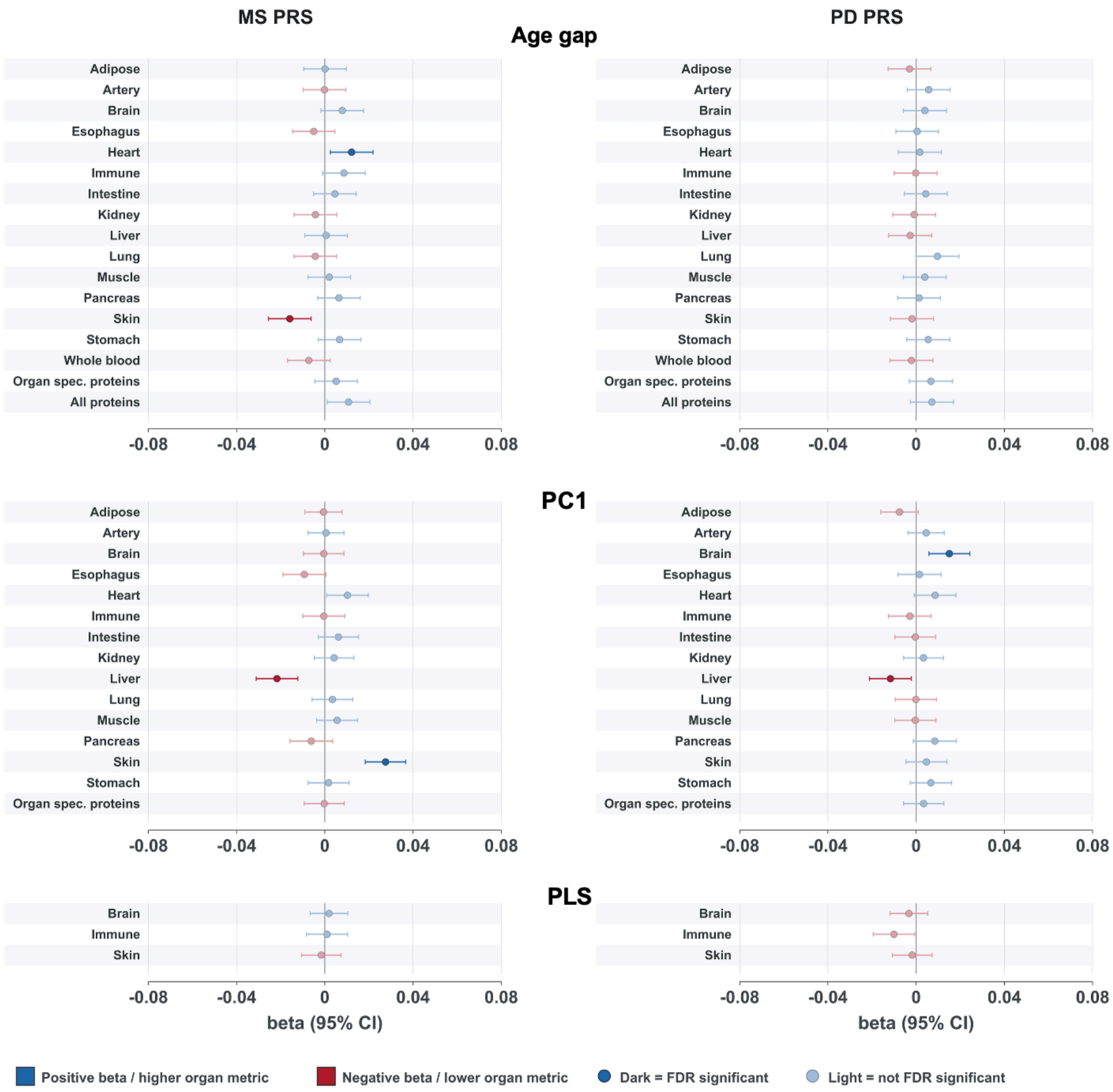


**Supplementary Figure 6. Associations between multiple sclerosis and Parkinson's disease polygenic risk scores and organ level proteomic metrics.**

Forest plots show associations between multiple sclerosis (MS) and Parkinson's disease (PD) polygenic risk scores and organ level proteomic metrics from multivariable linear models. Results are shown separately for age gap, PC1, and PLS1 metrics. Points mark beta estimates, and horizontal bars show 95% confidence intervals. All panels use the same x-axis range to support comparison. Blue indicates positive associations with higher metric values, and red presents negative associations with lower metric values. Darker points indicate FDR-significant associations, while lighter points indicate associations that did not survive FDR correction.

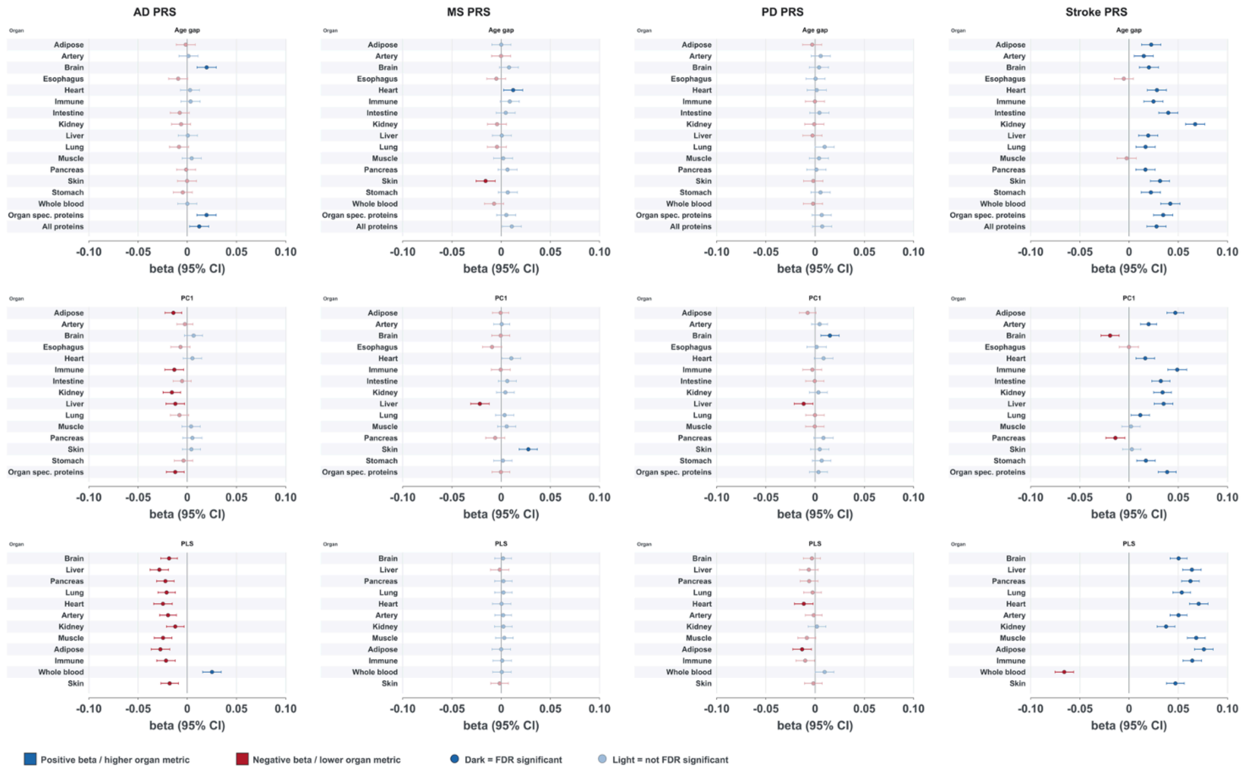


**Supplementary Figure 7. Associations between disease polygenic risk scores and organ level proteomic metrics.** Forest plots show associations between standardized polygenic risk scores for Alzheimer's disease, multiple sclerosis, Parkinson's disease, and stroke and organ level proteomic metrics. Models were fitted separately by disease PRS and metric class, with standardized PRS as the outcome and organ level proteomic metrics as predictors, adjusting for age and sex. Results are shown separately for age gap, PC1, and PLS1 metrics. Points indicate beta estimates and horizontal bars indicate 95% confidence intervals. Blue indicates positive associations between higher organ level proteomic metric values and higher PRS, while red indicates negative associations. Darker points indicate FDR-significant associations, and lighter points indicate associations that did not survive FDR correction.
